# A systematic, unbiased mapping of CD8^+^ and CD4^+^ T cell epitopes in Yellow Fever vaccinees

**DOI:** 10.1101/2020.03.28.012468

**Authors:** Anette Stryhn, Michael Kongsgaard, Michael Rasmussen, Mikkel Nors Harndahl, Thomas Østerbye, Maria Rosaria Bassi, Søren Thybo, Mette Gabriel, Morten Bagge Hansen, Morten Nielsen, Jan Pravsgaard Christensen, Allan Randrup Thomsen, Soren Buus

## Abstract

Examining CD8^+^ and CD4^+^ T cell responses after primary Yellow Fever vaccination in a cohort of 210 volunteers, we have identified and tetramer-validated 92 CD8^+^ and 50 CD4^+^ T cell epitopes, many inducing strong and prevalent (i.e. immunodominant) T cell responses. Restricted by 40 and 14 HLA-class I and II allotypes, respectively, these responses have wide population coverage and might be of considerable academic, diagnostic and therapeutic interest. The broad coverage of epitopes and HLA overcame the otherwise confounding effects of HLA diversity and non-HLA background providing the first evidence of T cell immunodomination in humans. Also, double-staining of CD4^+^ T cells with tetramers representing the same HLA-binding core, albeit with different flanking regions, demonstrated an extensive diversification of the specificities of many CD4^+^ T cell responses. We suggest that this could reduce the risk of pathogen escape, and that multi-tetramer staining is required to reveal the true magnitude and diversity of CD4^+^ T cell responses. Our T cell epitope discovery approach uses a combination of 1) overlapping peptides representing the entire Yellow Fever virus proteome to search for peptides containing CD4^+^ and/or CD8^+^ T cell epitopes, 2) predictors of peptide-HLA binding to suggest epitopes and their restricting HLA allotypes, 3) generation of peptide-HLA tetramers to identify T cell epitopes, and 4) analysis of *ex vivo* T cell responses to validate the same. This approach is systematic, exhaustive, and can be done in any individual of any HLA haplotype. It is all-inclusive in the sense that it includes all protein antigens and peptide epitopes, and encompasses both CD4^+^ and CD8^+^ T cell epitopes. It is efficient and, importantly, reduces the false discovery rate. The unbiased nature of the T cell epitope discovery approach presented here should support the refinement of future peptide-HLA class I and II predictors and tetramer technologies, which eventually should cover all HLA class I and II isotypes. We believe that future investigations of emerging pathogens (e.g. SARS-CoV-2) should include population-wide T cell epitope discovery using blood samples from patients, convalescents and/or long-term survivors, who might all hold important information on T cell epitopes and responses.

## 2. Introduction

The immune system can protect its host against virtually any invading pathogen; yet, it can also cause serious pathology. The ability to discriminate between foreign and self is key to exerting immune protection without inflicting immune pathology. Immune recognition is therefore of immense interest and efficient methods to identify and validate immune epitopes are a high priority. In this context, T cells, which effectively orchestrate the overall immune response, are of particular interest. T cells are specific for compound ligands consisting of peptides, generated intracellularly by proteolytic degradation of protein antigens, which are presented in the context of major histocompatibility complex (MHC) (or human leucocyte antigens (HLA)) molecules on the surface of antigen presenting cells (APC) (1). The interaction between peptide and HLA is specific; the resulting HLA-mediated T cell epitope selection process being greatly diversified by the polygenic and polymorphic nature of the HLA. This significantly affects the peptide-binding specificity of the set of HLA molecules that are available to any given host; something that effectively individualizes our immune responses. Although other events are also involved in antigen processing and presentation, the single most selective event is that of peptide-HLA binding. It is estimated that ≈0.5% of all possible peptide-HLA combinations are of a sufficiently high affinity that they potentially, but not necessarily, could be immunogenic (2). Major efforts have been devoted to understand, quantitate and preferably predict peptide-HLA binding as a means to identify T cell epitopes. Proposed in 1999, the “human MHC project” aims at mapping all human MHC (or HLA) specificities (3, 4). Established in 2004, the “Immune Epitope Database” (IEDB) has become an authoritative repository of HLA binding peptides and T cell epitopes, and of methods to predict these (5). The recent breakthrough in cancer immunotherapy has reinforced the interest in fast and efficient methods to identify T cell epitopes with special emphasis on identifying immunogenic neoepitopes for personalized cancer immunotherapy. Thus, several recent international research efforts, such as the “Human ImmunoPeptidome Project and Consortium”, “Tumor Neoantigen Selection Alliance” and others, have focused on T cell epitope discovery. Employing recent advances in mass spectrometry to perform large-scale identification of peptides eluted of HLA molecules, these efforts promise to identify natural ligands thereby capturing information on both antigen processing and HLA binding (6).

Over the past decades, substantial progress has been made on predicting peptide-HLA interactions, particularly for HLA class I (HLA-I), which restricts CD8^+^ cytotoxic T cells (CTL’s), and to a lesser degree on predictions for HLA class II (HLA-II), which restricts CD4^+^ helper T cells (Th) (7-15). State-of-the-art predictors such as NetMHCpan, an artificial neural network method based on a large collection of experimental peptide-HLA-I binding data, can successfully identify 96.5% of CD8^+^ T cell epitopes, while rejecting 98.5% of non-epitopes (16). However, considering that only 1 of 2000 (2) to 8000 (17) random peptides is a T cell immunogen in the context of a given HLA molecule, even a rejection rate as high as 98.5% translates into a high false discovery rate (FDR) (8, 10, 11, 18). This is a general problem of current peptide-HLA binding predictors (10, 11), and it is particularly problematic when trying to develop a neoepitope-specific, personalized cancer immunotherapy where timely delivery of a few unique cancer neoepitopes is of paramount importance; something that potentially could be achieved with even better predictors (8, 19-21).

Yellow Fever Virus (YFV) is a mosquito-borne flavivirus (i.e. a ssRNA virus) (22, 23). It remains an important human pathogen despite the existence of an effective live attenuated vaccine (24). Particularly relevant to this study, previous analyses of the CD8^+^ T cell response against a limited number of epitopes have revealed that vaccination with this live vaccine represents an excellent model for studying the host response to a viral infection (25, 26). The main advantages are that the precise time and the exact identity of the immune challenge are both known (note that the vaccine strain used here is known to be stable (27)); issues that otherwise might complicate the interpretation of immune responses observed in patients that are naturally infected with a variable pathogen. Here, we have generated a comprehensive, population-wide T cell epitope discovery approach with a much-reduced FDR, and used it to identify and validate immunodominant CD8^+^ and CD4^+^ T cell epitopes in a cohort of 210 HLA-typed, primary YFV vaccinees. This involves using a “forward (or direct) immunology” approach, where you start with a specific T cell response of interest and then search for the epitope(s) being recognized (28, 29), to perform an initial identification of T cell stimulatory peptides. Subsequently, a “reverse immunology” approach, where you start by predicting possible T cell epitopes and then search for a T cell response of the corresponding specificity (30, 31) was used, to perform a final identification and validation of the underlying specific T cell epitopes and their HLA restriction elements. From hereon, this approach is denoted as a “hybrid forward-reverse immunology” (HFRI) approach. Briefly, in the “forward immunology” step, PBMC’s obtained 2-3 weeks after primary YFV vaccination were *ex vivo* stimulated with an overlapping peptide library representing the entire 3411 amino acid YFV proteome and tested by an IFNγ-specific intracellular cytokine secretion (ICS) assay thereby identifying CD8^+^ and CD4^+^ T cell stimulatory YFV-derived peptides. In the subsequent “reverse immunology” step, predictors were used to select appropriate peptide-HLA combinations for the generation of peptide-HLA tetramers, which then were used to identify and validate the underlying T cell epitopes and their HLA restriction elements. Applying this HFRI approach to T cell epitope discovery in 50 YFV vaccinees, we identified and tetramer-validated 92 CD8^+^ and 50 CD4^+^ T cell epitopes covering 40 HLA-I and 14 HLA-II allotypes, respectively (note that he tetramer-validation step could not be performed exhaustively for the CD4^+^ T cell epitope discovery process and that the true number of CD4^+^ T cell epitopes probably was many times larger than the 50 validated CD4^+^ T cell epitopes reported here). With a cohort of 210 YFV vaccinees, the prevalence of responses against the CD8^+^ T cell epitopes could be examined. About half (45%) of these epitopes were recognized in >90% of the individuals expressing the HLA-I in question. By this token, they could be considered strongly immunodominant. We conclude that T cell epitope discovery using this HFRI approach is highly efficient, in particular when examining larger populations responding to the same pathogen (e.g. an infectious pathogen e.g. SARS, Ebola, Zika, SARS-CoV-2). Furthermore, we suggest that the HFRI approach is unbiased and that the resulting T cell epitopes should serve as a valuable benchmark for future improvements of predictive algorithms of immunogenicity.

## 3. Results

### 3.1. Obtaining blood samples from HLA-typed yellow fever vaccinees

Primary vaccination with the attenuated YFV vaccine, 17D-204, is known to trigger a prompt and vigorous cellular immune reaction (25, 26). Here, 210 vaccinees were recruited, and peripheral blood mononuclear cells (PBMC) were prepared from 50- and 200-ml blood samples obtained before and ≈2 weeks after primary vaccination, respectively (26). The typical yield from the latter was ≈ 450 million PBMC. All vaccinees were HLA typed at high-resolution (i.e. 4 digit) including all nine classical, polymorphic HLA loci (i.e. HLA-A, B, C, DRB1, DRB3/4/5, DQA1, DQB1, DPA1 and DPB1) (26).

### 3.2. Overlapping peptides representing the entire yellow fever virus proteome

The 17D-204 vaccine encodes a single polyprotein precursor of 3411 amino acids (aa), which is processed into 15 proteins. The full genome (GenBank accession# X15062) and proteome (Swiss-Prot accession# P03314) sequences of the 17D-204 have been determined (32). A library of 850 overlapping 15mer peptides overlapping by 11 aa, spanning the entire YFV precursor protein (essentially the YFV proteome), was generated. Additionally, 50 peptides representing potentially aberrant YFV translation products were selected. Of the resulting 900 peptides, synthesis failed for 30 peptides (3%) leaving 870 peptides for analysis.

### 3.3. Matrix-based screening strategies

Since testing each of these peptides individually would exhaust the available PBMC’s, the peptides were tested in pools. Initially, the peptides were organized into a single 30×30 matrix from which 30 “column pools” and 30 “row pools” were generated leading to a total of 60 pools each containing ≈30 different peptides. Each peptide would be present in two pools: one column and one row pool (**Supplementary Figure S3**). The intersections of stimulatory column and row pools should ideally identify which peptide might be immunogenic and therefore should be further investigated on an individual basis.

This 30×30 matrix strategy was initially tested using an *ex-vivo* IFNγ ELISpot assay as readout. After the first 94 primary vaccinated donors had been recruited, the average number of positive column/row intersections was found to be 418 (range 26 to 870) (**Figure 1A**) suggesting that the hit rate from pools containing 30 peptides was too high, at least in the setting of this acute viral response, to be effective in eliminating non-stimulatory peptides from further consideration.

**Figure 1:**
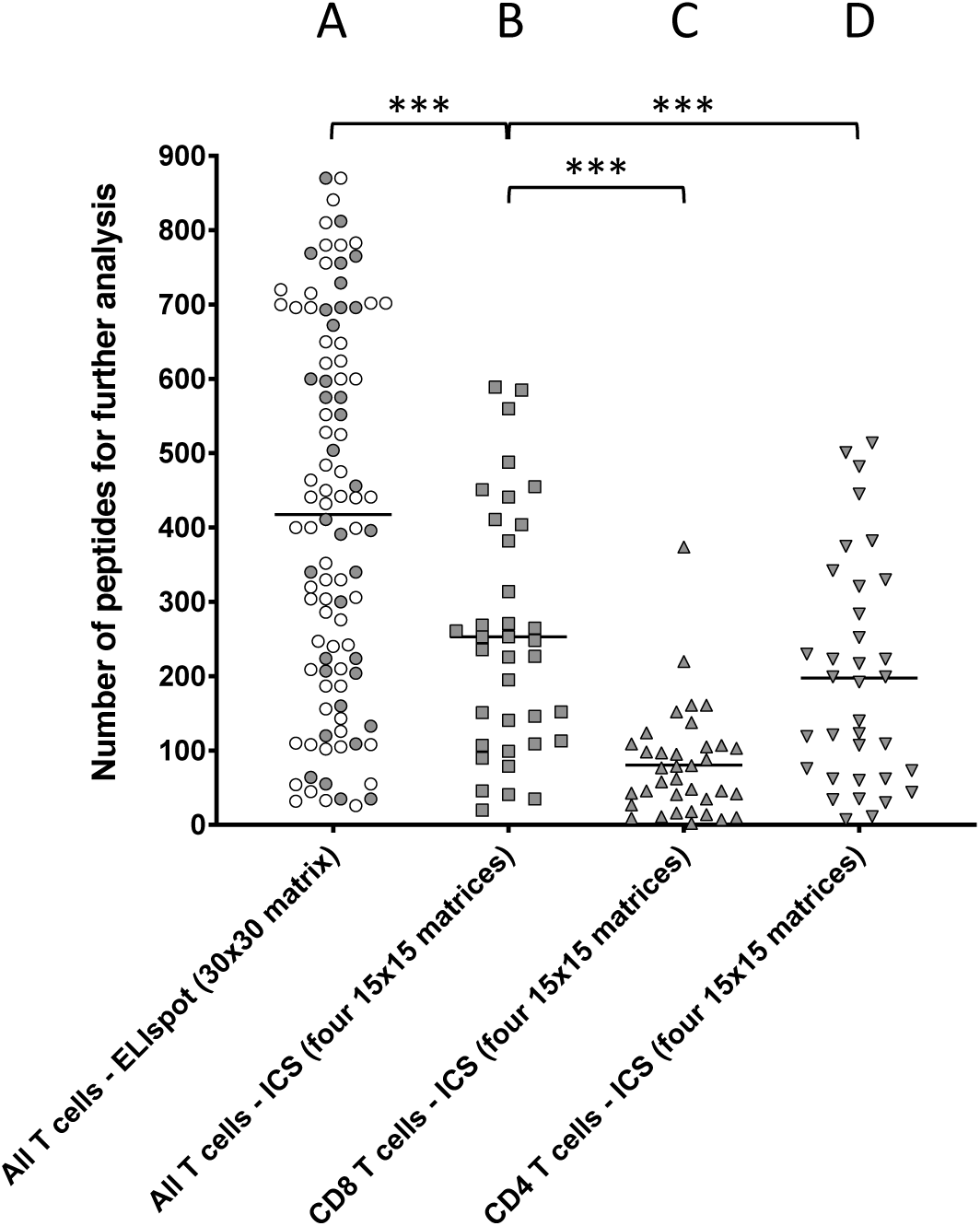
Comparing the number of peptide-containing peptides identified by the two different approaches. T cells obtained *ex vivo* from primary YFV vaccinees were stimulated with matrix-derived pools of YFV peptides and responses were read by IFNg-specific ELIspot or ICS. The peptides were distributed into matrixes, column and row pools of peptides were generated, and used to test T cell stimulation. Intersections of stimulatory column and row pools putatively identified single stimulatory peptides for further analysis (**Supplementary Figure S3**). A) Peptides were distributed into one 30×30 matrix generating 30+30 = 60 pools, which were used to stimulate T cell responses in 94 donors using an IFNg-specific ELISpot assay as readout of all (i.e. CD4^+^ and CD8^+^) T cell responses (average 418 positive intersections (range 26–870)). B-C) Peptides were distributed into four ≈15×15 matrices generating 4x(≈15+15) ≈120 pools, which were used to stimulate T cell responses in 36 donors using an IFNg-specific ICS assay as readout of B) all T cell responses (average 253 intersections (range 20–589)), C) CD8^+^ T cell responses (average 80 intersections (range 2–374), and D) CD4^+^ T cell responses (average 197 intersections (range 7–514). The symbols representing the 36 donors that were examined by both ELISpot and ICS have been shaded. Mann Whitney U test was used to determine the significance of the difference between the indicated groups (***p□<□0.0001).

To reduce the hit rate per peptide pool, the peptides were re-organized into four smaller matrices, three 15×15 matrices and one 14×15. For each matrix, 14 to 15 column pools and 15 row pools were generated leading to a total of 119 pools, which each contained 14 to 15 different peptides (**Supplementary Figure S3)**. To further reduce the number of relevant intersections, the IFNγ ELISpot assay was replaced by an IFNγ intracellular cytokine staining (ICS) assay, which can discriminate between CD8^+^ and CD4^+^ T cells and therefore eliminate intersections with mismatched CD8^+^ and CD4^+^ T cell responses. Furthermore, to increase the number of T cells available for the ICS assay, PBMC were expanded in four separate *in vitro* cultures containing a pool of ≈225 peptides corresponding to each of the four matrices, respectively. After 8 days, each matrix-expanded PBMC culture was tested against the appropriate row and column pools using IFNγ ICS as readout. For comparison, 36 donors, which had already been analyzed using the 30×30-matrix, ELISpot-based screening strategy, were re-screened using the 4x(15×15)-matrix, ICS-based screening strategy (**Figure 1B-C**). The aggregated CD4^+^ and CD8^+^ T cell responses were calculated for ICS responses (denoted “All T cells” in Figure 1) and compared those from Elispot reponses. The total number of intersections needing deconvolution was significantly lower for the 4×(15×15 ICS strategy (average intersections 253 (range 20–589)) than for the 30×30 ELISpot strategy (average 418 (range 26–870), (p<0.0001, N=36, Mann Whitney U test), **Figure 1A vs B**). The 253 intersections, which on average were detected by the ICS-based screening strategy, could further be broken down into an average of 80 (range 2-374) (**Figure 1C)** and 197 (range 7 – 514) (**Figure 1D**) intersections representing CD8^+^ and CD4^+^ T cell responses, respectively (30 of these intersections were shared). The peptides corresponding to these intersections were subsequently tested individually to identify which of the intersections truly represented CD8^+^ and/or CD4^+^ T responses.

### 3.4. Identification of stimulatory 15mer peptides (exemplified by donor YF1067)

The complete screening and validation procedure is illustrated using donor YF1067. The blood sample for donor YF1067 was collected at day 16 post vaccination, which is within the time span of optimal post vaccination YFV responses (25, 26). It gave a relatively high yield of 700*10^6^ PBMC’s for the subsequent epitope discovery effort. Donor YF1067 was initially analyzed by the 30×30-matrix, IFNγ ELISpot-based screening strategy where 690 positive intersections were identified. A total (i.e. cumulative) YFV-specific response of 8000 SFU were obtained suggesting a T cell response of considerable breath and magnitude. Re-analyzing this donor using the four-matrix, ICS-based screening strategy, the number of intersections for follow-up analysis could be reduced to 253; 78 representing CD8^+^ T cell responses and 218 representing CD4^+^ T cell responses (43 of the intersections contained both CD4^+^ and CD8^+^ T cell responses). Peptides corresponding to the 253 intersections were tested individually by ICS; this identified 27 and 31 CD8^+^ and CD4^+^ T cell stimulatory 15mer peptides, respectively (**Table I and II**). The next steps aimed at identifying the underlying CD8^+^ and CD4^+^ T cell epitope(s) and their HLA restriction element(s), preferably by generating the corresponding tetramer(s), and validate the epitope(s). For a general outlined of this epitope discovery scheme, see **Figure 2**.

**Table I:**
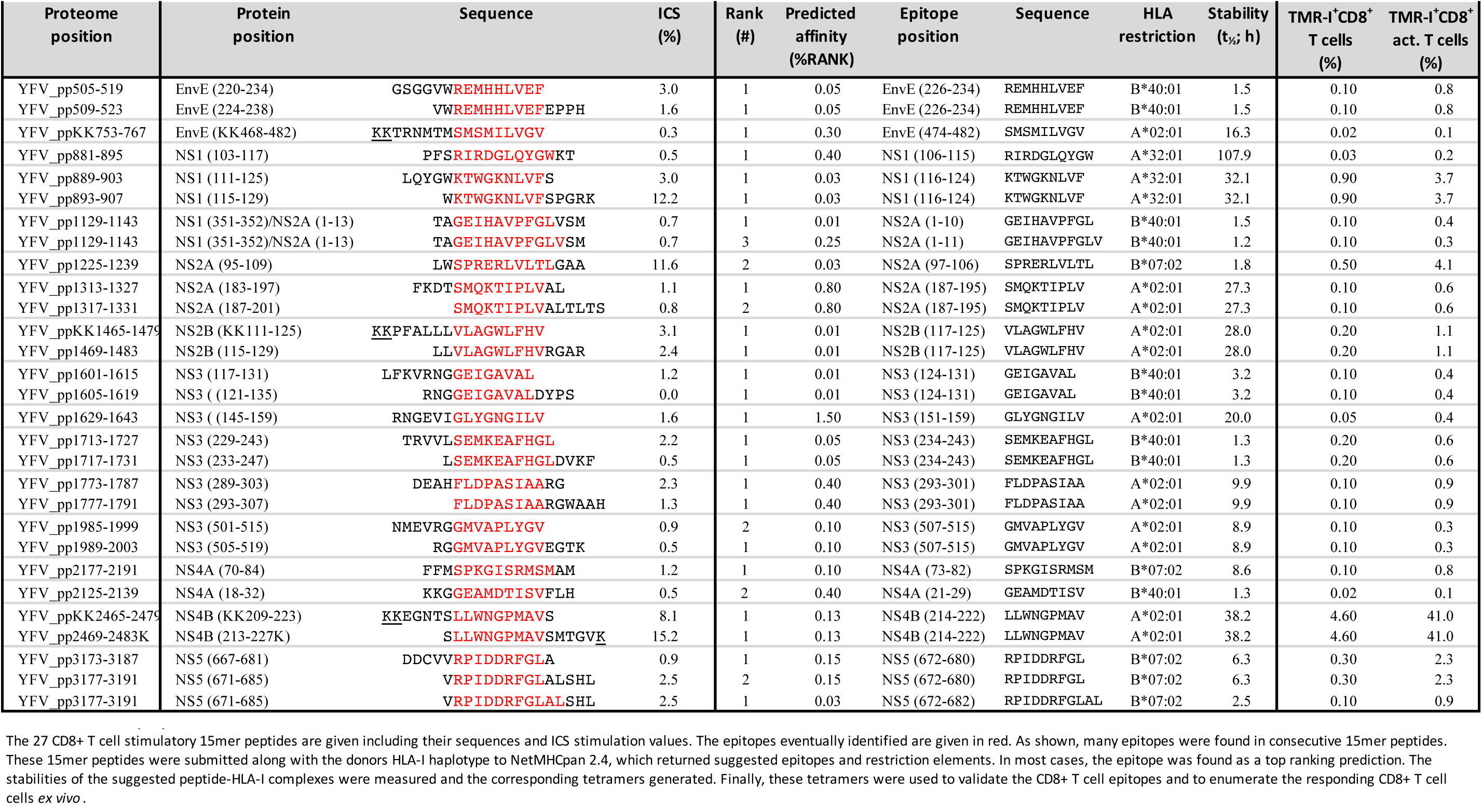
CD8+ T cell epitopes identified in donor YF1067.

**Table II:**
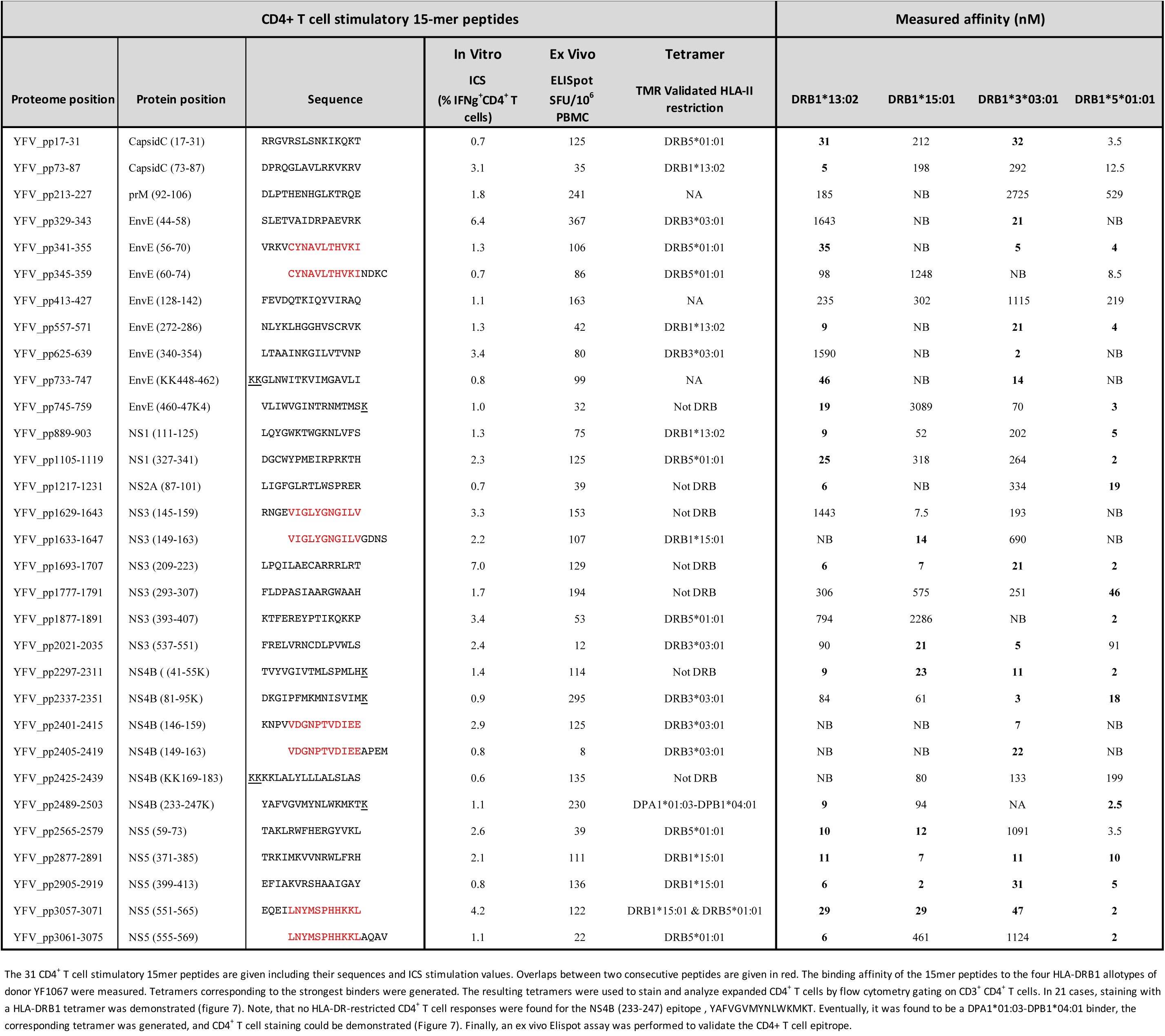
CD4+ T cell epitopes identified in donor YF1067.

**Figure 2:**
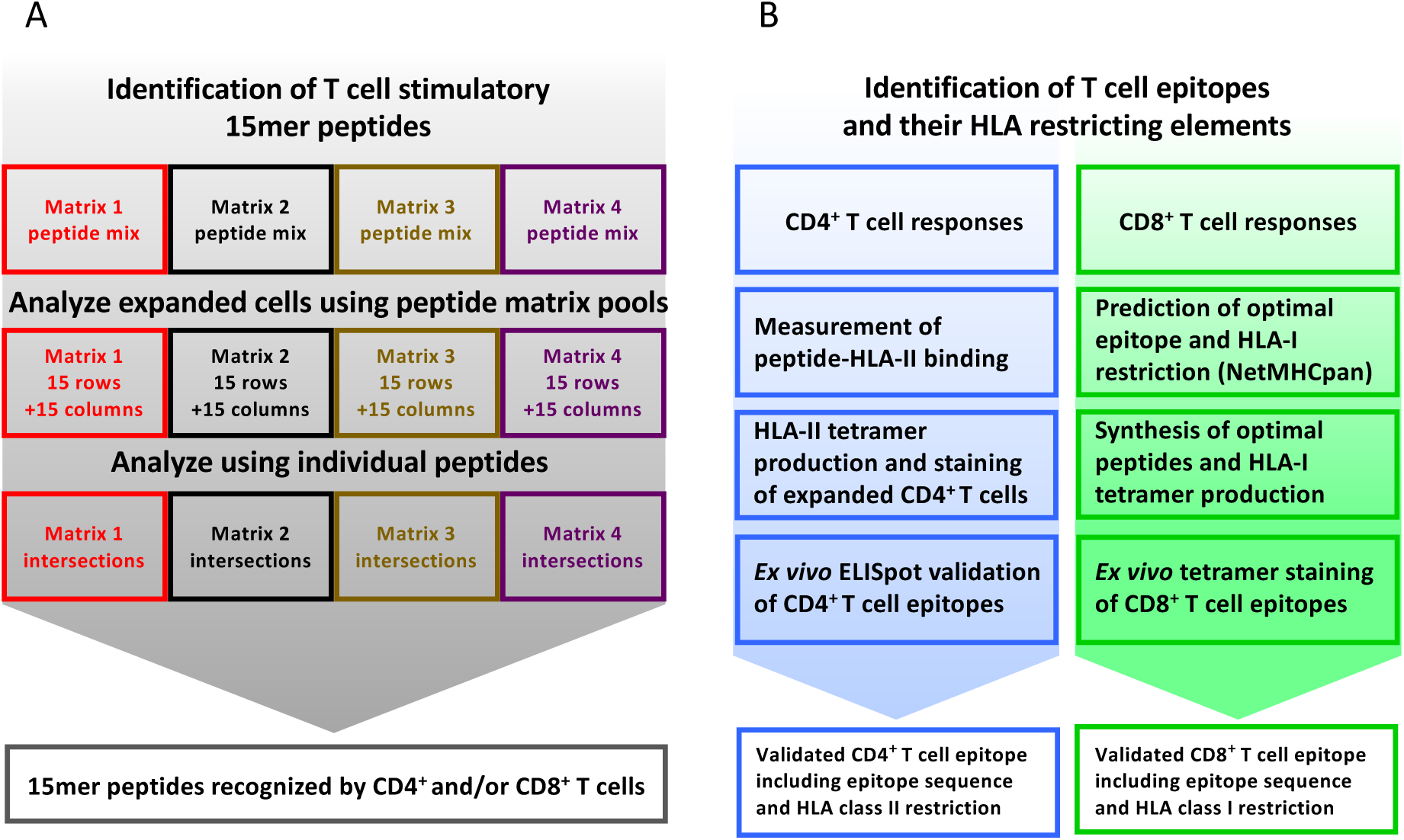
T cell epitope screening strategy. **A) Identification of stimulatory 15mer peptides:** PBMC’s from YFV vaccinated donors were divided into four cultures and in vitro stimulated with peptide sublibraries corresponding to each of the four 15×15 peptide matrices. After 8 days, each sublibrary expanded PBMC culture was tested by ICS against the matrix-specific row and column peptide pools. Subsequently, individual peptides representing stimulatory matrix intersections were analyzed to identify single T cell stimulatory 15-mer peptides. **B) Identification of T cell epitopes and their HLA restriction elements:** *CD4*^*+*^ *T cell epitope deconvolution*: Single CD4^+^ T cell stimulatory 15mer peptides were tested for binding to the donor’s HLA-DR molecules using a biochemical HLA class II binding assay, positive interactions were used to generate peptide-HLA class II tetramers, and these tetramers were used to stain expanded T cells, and the resulting epitopes were eventually validated by ex vivo ELISpot analysis. *CD8*^*+*^ *T cell epitope deconvolution*: Single CD8^+^ T cell stimulatory 15mer peptides were submitted to the NetMHCpan 2.4 predictor together with the donor’s HLA class I haplotype to identify optimal epitopes and their HLA-restriction elements. These optimal epitopes were subsequently synthesized and validated by ex vivo peptide-HLA class I tetramer staining.

### 3.5. CD8^+^ T cell epitope discovery

#### 3.5.1. Identification and validation of CD8^+^ T cell epitopes exemplified by donor YF1067

The sequences of the twenty-seven 15mer peptides, which stimulated CD8^+^ T cell responses in donor YF1067, were submitted, along with the donor’s HLA-I allotypes (*in casu* HLA-A*02:01, -A*32:01, -B*07:02, -B*40:01, -C*03:04 and -C*07:02), to our webserver NetMHCpan (version 2.4 at http://www.cbs.dtu.dk/services/NetMHCpan-2.4 was available at the time of this analysis). *In silico*, this predictor considered all 26 submer peptides of 8-11mer length, which could possibly be generated from a 15mer peptide, and predicted their binding to all six HLA-I allotypes of donor YF1067, a total of 6*26 = 156 submer-HLA-I combinations per 15mer peptide, and returned a ranked list across all six HLA-I allotypes of the most likely epitope(s) and their HLA-I restriction element(s). For all 27 CD8^+^ T cell stimulatory peptides, this amounted to predicting the binding affinities of 27*156 = 4212 submer-HLA-I combinations. For each 15mer peptide, submers representing the top one to three predicted affinities were synthesized and the stabilities of the corresponding peptide-HLA-I interactions were measured experimentally (**Table I**). Fluorochrome-labeled tetramers corresponding to the most stable peptide-HLA-I interactions were generated and used to label relevant CD8^+^ T cells. When available, surplus T cells from the initial expansion cultures were used as a first line of identification of CD8^+^ T cell epitopes and their restriction elements, however, *ex vivo* tests were always used for the final CD8^+^ T cell epitope validation, and for enumerating and characterizing epitope-specific CD8^+^ T cells. The matrix-identified CD8^+^ T cell stimulatory 15mer peptides and the corresponding tetramer-validated optimal CD8^+^ T cell epitopes and their restriction elements are listed (**Table I**). For each of the 27 CD8^+^ T cell stimulatory 15mer peptides identified in donor YF1067, one or more CD8^+^ T cell epitopes and their HLA-I restriction elements were identified. Some of the epitopes were present in two consecutive overlapping 15mer peptides and should therefore only be counted as epitopes once. With this in mind, 19 unique CD8^+^ T cell epitopes were recognized by donor YF1067 (7 epitopes restricted by HLA-A*02:01, 2 by HLA-A*32:01, 4 by HLA-B*07:02, 6 by HLA-B*40:01, and none by HLA-C*03:04 or -C*07:02) (**Table I**).

CD8^+^ T cell specific for the 19 unique YFV-derived epitopes were readily detectable and enumerable *ex vivo* during the acute primary response of donor YF1067. The frequencies of total, as well as activated, tetramer-positive CD8^+^ T cells were determined (**Table I**). The most frequently and immunodominant epitope of all, the HLA-A*02:01-restricted NS4B_214-222_ epitope, was recognized by 4.6% of CD8^+^ T cells in donor YF1067. The frequencies of CD8 T cells recognizing each of the other 18 epitopes ranged from 0.03-0.9%; the total frequency of CD8^+^ T cells recognizing the 19 YFV epitopes was ≈8%. The YFV vaccine induced a measurable increase in the overall frequency of activated CD8^+^ T cells (i.e. CD38^+^HLA-DR^+^CD8^+^ T cells) (26). In donor YF1067, the YFV vaccine induced an increase in activated CD8^+^ T cell from 0.6% pre- to 7% post-vaccination. Notably, the HLA-A*02:01-restricted NS4B_214-222_-epitope was recognized by 41% of the activated CD8^+^ T cells in donor YF1067. The frequencies of activated CD8^+^ T cells recognizing each of the other 18 epitopes ranged from 0.1-4.1%. In total, the 19 identified YFV-specific CD8^+^ T cell epitopes accounted for the majority (≈60%) of activated CD8^+^ T cells observed during the acute response following primary YFV vaccination.

#### 3.5.2. Extending CD8^+^ T cell epitope discovery to 50 primary YFV vaccinated individuals

The CD8^+^ T cell epitope discovery strategy described above for donor YF1067 was extended to 50 randomly selected donors, who were sampled at day 12-21 after vaccination i.e. at the peak of a primary anti-YF vaccine response (25, 26). CD8^+^ T cell responses specific for 120 different peptide HLA-I combinations were identified and validated by *ex vivo* tetramer staining (for an overview, see **Figure 3**, and for details, see **Supplementary Table SI**). This represented 92 different CD8^+^ T cell epitopes restricted by 40 different HLA-I molecules; 68, 20 and 4 epitopes were restricted by 1, 2 and 3 different HLA-I molecules, respectively. The HLA-A, -B and -C allotypes covered by the 50 donors were respectively 19, 30 and 20 of which the majority, 15, 27 and 16, were available to us for tetramer validation. Thirteen of the 15 different HLA-A allotypes tested served as restriction elements of 38 different CD8^+^ T cell peptide epitopes leading to the presentation of 44 immunogenic peptide-HLA-A combinations; 26 of the 27 different HLA-B allotypes tested served as restriction elements of 56 different epitopes leading to the presentation of 74 immunogenic peptide-HLA-B combinations; whereas only one of the 16 different HLA-C allotypes tested served as restriction elements of 2 different epitopes leading to the presentation of 2 immunogenic peptide-HLA-C combinations. The average number of CD8^+^ T cell epitopes identified per HLA-A and -B allotype, 3.4 and 2.8, respectively, were not significantly different (P>50%, Fishers exact test, two-tailed (GraphPad)).

**Figure 3:**
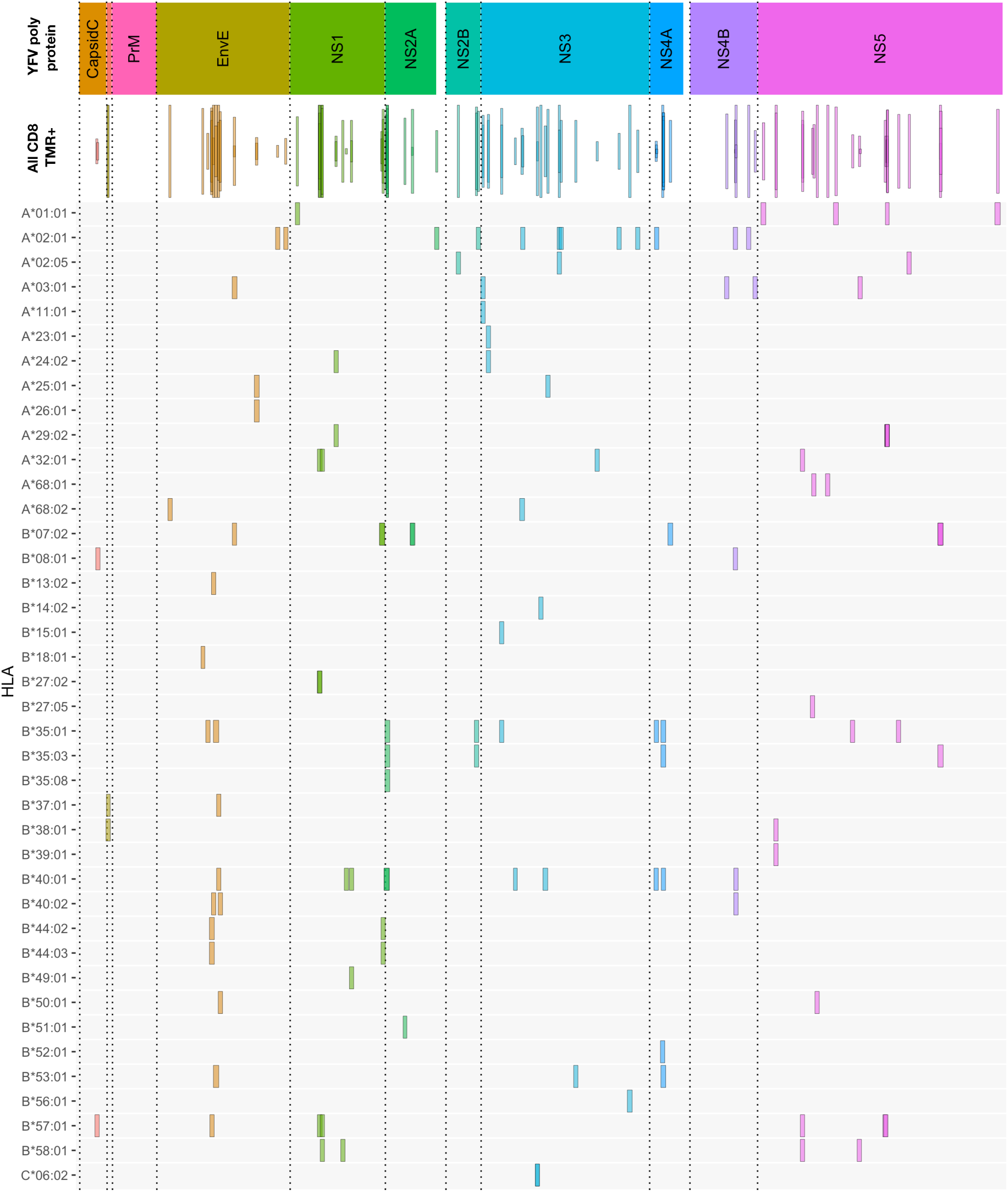
Overview of CD8^+^ T cell epitopes discovered in 50 primary YFV vaccinated donors. The upper bar indicates the YFV polyprotein. The individual proteins have been color-coded, and stippled vertical lines are used as further guidance to delineate each protein throughout the rest of the figure. The “all CD8” bar indicates the positions of each CD8^+^ T cell stimulatory 15mer peptide relative to the YFV polyprotein. Each 15mer is shown as a frame that horizontally indicates the starting and ending positions of the 15mer peptide, vertically indicates the number of donors, of the 50 donors tested, who responded to the peptide, and the color-coded is according to the polyprotein coloring scheme (to enhance the visualization of overlapping peptide sequences, this coloring is translucent). The lower HLA-I allotype-designated bars indicate the tetramer-validated epitopes and their HLA-I restriction elements (e.g. A*01:01 is shorthand for HLA-A*01:01). Again, the frame horizontally indicates the starting and ending positions of the epitopes; however, for visual clarity, all frames have the same vertical dimension. The details of each epitop (epitope sequence, CD8+ T cell stimulation, ex vivo tetramer staining frequency, and response prevalence is given in **Supplementary table SI**)

To the best of our knowledge, 84 of the 92 YF-specific CD8^+^ T cell epitopes, and 110 of the 120 epitope-HLA-I combinations reported here and in previous publications (26, 33), were first identified as a result of this HFRI project. For the previously reported epitopes or epitope-HLA-I combinations, minor adjustments of the already available information could be made: some had not been tetramer validated before, and others were also found to be restricted by other, albeit closely related, HLA-I allotypes than those previously reported. In a few cases, tetramers representing the exact epitope-HLA-I combinations previously reported failed to label CD8^+^ T cells in our donors despite expressing the appropriate HLA-I allotype (for details see **Supplementary Table SI**). Our in-house peptide repository included 533 YFV-derived peptides from previous HLA mapping efforts (34). Using the contemporary NetMHCpan2.4 at %Rank cut-off of 0.5% to select putative binders from this repository, we generated 90 additional peptide-HLA-I tetramers (i.e. tetramers that had not already been prepared in the course of the present HFRI approach). We included these tetramers in the immunodominance analysis described below. Nine additional peptide-HLA-I combinations, which had not been observed previously, were identified; four representing previously identified epitope presented by an alternative HLA-I restriction element, and five representing new YFV-specific CD8^+^ T cell epitopes (**Supplementary Table SII).** Thus, the total number of CD8^+^ T cell epitopes discovered and tetramer validated here was 97 of which 92 (or 95%) were identified by the HFRI approach.

#### 3.5.3. Extending CD8^+^ T cell epitope discovery to additional donors to address immunodominance

We systematically extended the analysis of *ex vivo* responses to additional donors expressing relevant HLA-I restriction elements and evaluated them in terms of *prevalence* (the frequency of responders in donors with the HLA-I restriction element in question) and *response magnitude* (the average *ex vivo* frequency of tetramer positive, activated CD8^+^ T cells of the responding donors) **Supplementary Table SI and SII**. To allow for a reasonable assessment of prevalence, the final analysis included epitopes restricted by HLA-I molecules represented by at least 5 donors, who had donated blood samples 12-21 days after vaccination. This involved a total of 98 peptide-HLA combination representing 81 epitopes presented by 24 HLA-I allotypes (**Figure 4**). Immunodominance was frequently observed. From an epitope point of view, 25 (or 31%) of the 81 epitopes had a prevalence of ≥90% and a median magnitude >0.03%, and 50 (or 62%) had a prevalence of ≥50% and a median magnitude >0.02%. From an HLA point of view, 16 (or 67%) of the 24 HLA-I molecules presented at least one epitope with ≥90% prevalence, and all 24 HLA-I molecules presented at least one epitope with at least 50% prevalence. In terms of HLA-I coverage and immunodominance, the vast majority of our cohort, 97%, 79% and 43%, carried at least one, two or three HLA-I allotypes, respectively, which presented at least one epitope with ≥90% prevalence. A selection of 10 immunodominant epitopes representing the most frequent HLA-A and -B allotypes would cover 95% of the Caucasian population.

**Figure 4:**
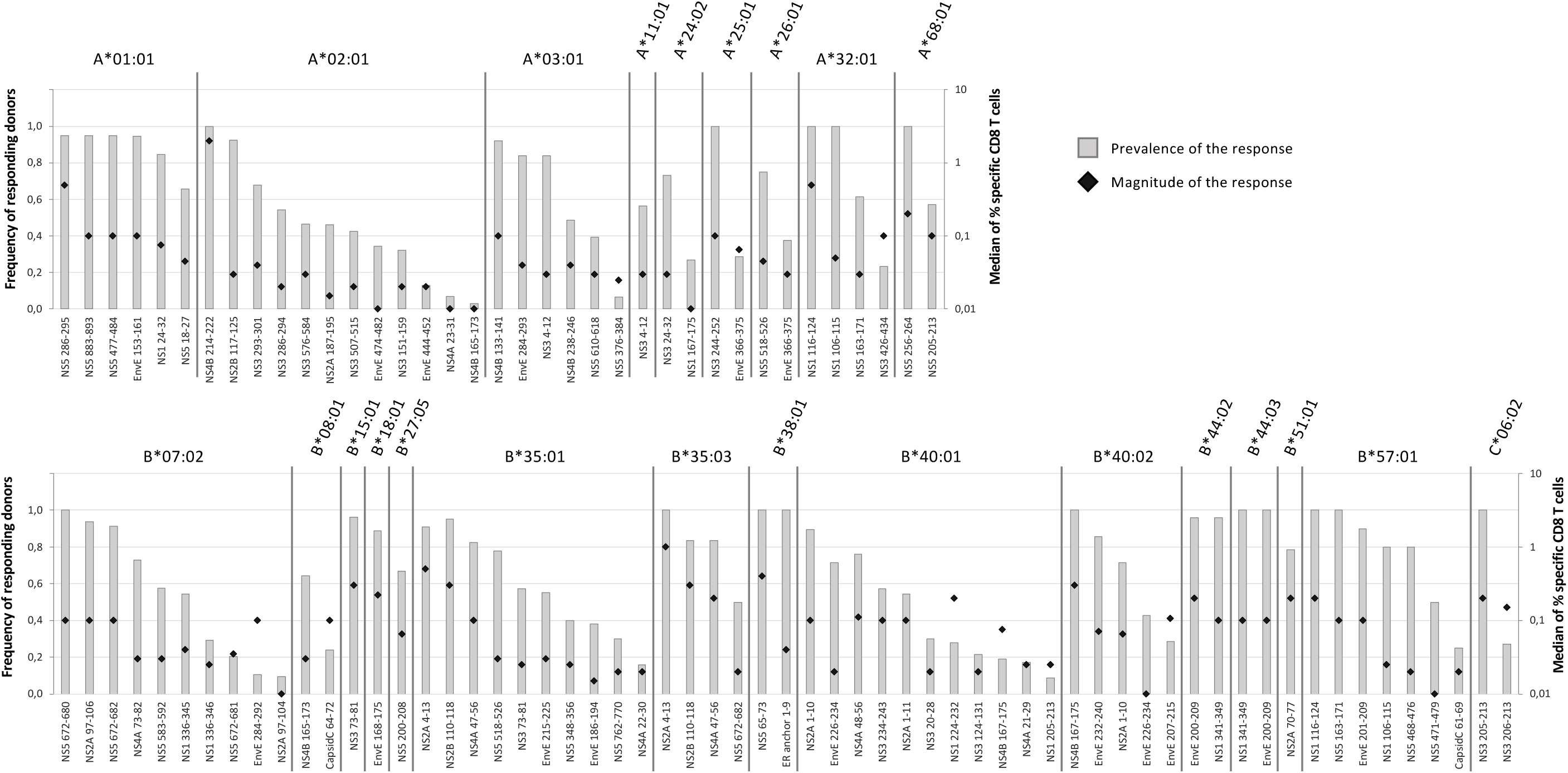
Prevalence and magnitude of CD8^+^ T cell responses. To determine the prevalence and the median magnitude of the CD8^+^ T cell responses towards the epitopes discovered in Table II, additional donors expressing the relevant HLA-I restriction elements were examined. The prevalence (grey columns) and the median magnitude of the responses (black diamond) were determined for each epitope-HLA combination. Only epitope-HLA combinations analyzed in 5 or more donors were included. The epitopes are organized according to restriction elements. The top figure shows the HLA-A restriction elements; the bottom figure shows the HLA-B and -C restriction elements.

#### 3.5.4. Inhibition of CD8^+^ T cell responses by immunodomination in an outbred human population

It has been suggested that immunodominant epitopes can curtail responses to other epitopes (reviewed in 35). The HLA-A*02:01 restricted, YFV NS4B_214-222_-epitope may represent a unique opportunity to address this in an outbred human population: it represents an exquisitely dominant CD8^+^ T cell response as all 93 HLA-A*02:01-positive donors examined here responded to this epitope and an average of 29% of all activated CD8^+^ T cells from *ex vivo* blood samples obtained 2-3 weeks after YFV vaccination were specific for this epitope. It has recently been suggested that this massive response can be explained by the invariant CDR1α loop of TRAV12-2 taking part in the recognition of this epitope (36). In donors, who had donated blood samples at the peak of the response (12-21 days after vaccination), we examined whether the presence of HLA-A*02:01, - A*01:01, or -A*03:01, could be correlated to the strength of CD8^+^ T cell responses restricted by other restriction elements, *in casu* all available HLA-B allotypes. We included 142 donors, which respectively could be split into 71 and 71 HLA-A*02:01 positives and negatives, 39 and 103 HLA-A*01:01 positives and negatives, or 30 and 112 HLA-A*03:01 positives and negatives; and used tetramers to examine the *ex vivo* frequencies of up to 45 different HLA-B-restricted responses. In the presence or absence of each of the three HLA-A restriction elements, the average frequencies of each of the HLA-B-restricted responses were determined leading to the generation of up to 45 matched-pairs per HLA-A. The frequencies, or magnitude, of the HLA-B-restricted responses were significantly reduced in the presence vs. absence of HLA-A*02:01 (median reduction of 0.032%, P<0.0001). In contrast, in the presence vs. absence of HLA-A*01:01, which have lesser immunodominant CD8^+^ T cell responses, there was a smaller and marginally significant reduction (median reduction of 0.008%, P = 0.09); in the presence vs. absence of HLA-A*03:01, which have even fewer immunodominant CD8^+^ T cell responses, there was a very small and non-significant increase (median increase of 0.003%, P = 0.54) (Wilcoxon Signed-Rank Test, **Figure 5**). We suggest that this may be the first demonstration of immunodomination in an outbred human population. Elucidating the underlying mechanisms is beyond the scope of this paper.

**Figure 5:**
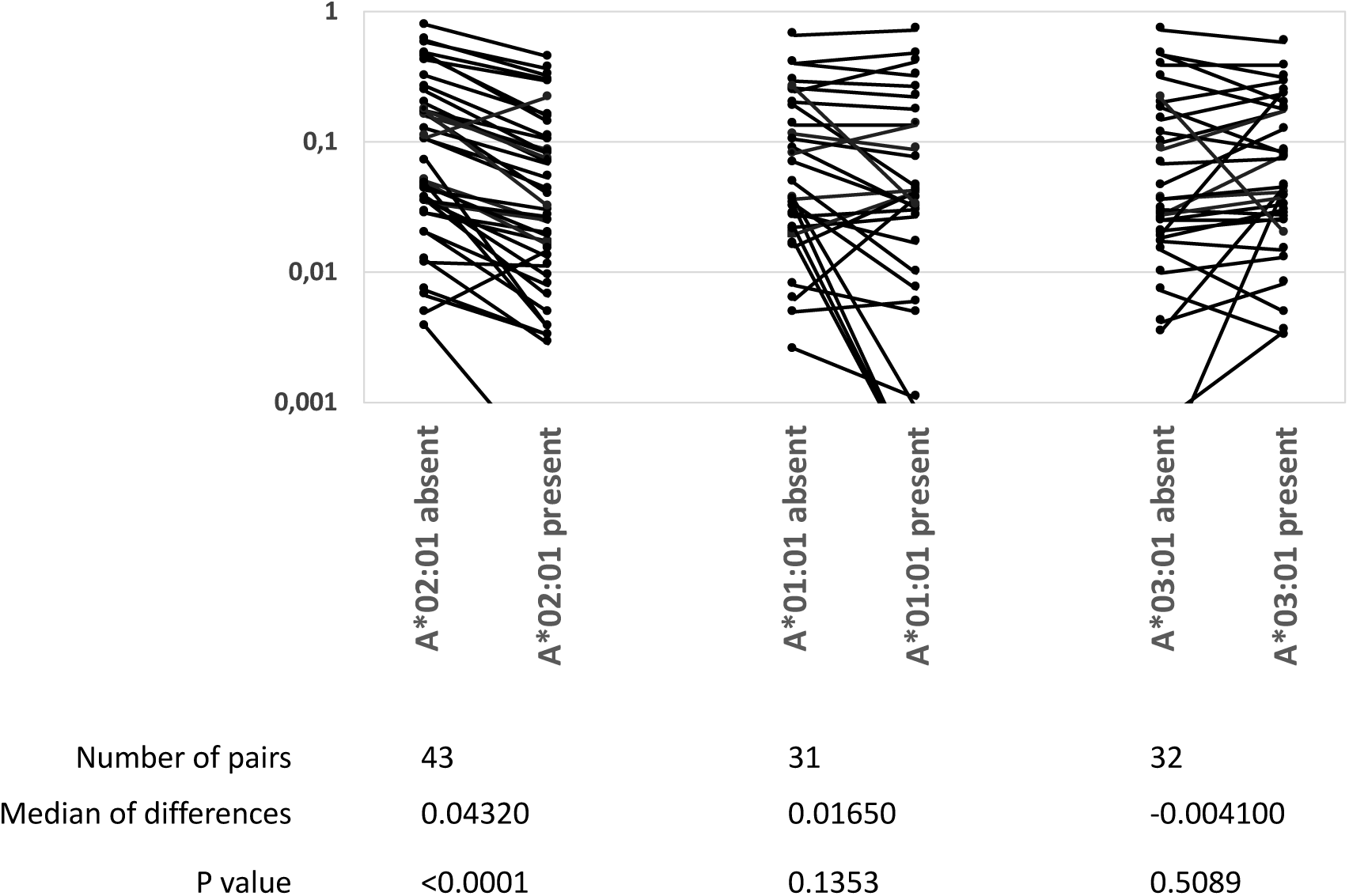
A highly immunodominant CD8^+^ T cell response correlates with a reduction of other CD8^+^ T cell responses. The NS4B_214-222_-specific, HLA-A*02:01-restricted CD8^+^ T cell response is highly immunodominant. The presence or absence of HLA-A*02:01, HLA-A*01:01 or HLA-A*03:01 in 142 donors were correlated with the magnitudes (measured as % TMR^+^, CD8^+^ T cells) of up to 45 YFV-specific, HLA-B-restricted CD8^+^ T cell responses. The data was analyzed by Wilcoxons matched-pair signed rank test (GraphPad Prism 8)

#### 3.5.5. CD8^+^ T cell epitope length distribution and recognition of size variants

The length of the 97 discovered CD8^+^ T cell epitopes ranged from 8mers to 11mers with a predominance of 9mers (67 (69%) 9mers, 18 (19%) 10mers, 5 (5%) 11mers and 7 (7%) 8mers) (**Supplementary Figure S1**). This matches well with available data for peptides eluted of HLA-A and -B molecules (33). Some of the epitopes were size variants of the same peptide sequence. In six cases, such size variants were presented by the same HLA-I restriction element (four cases involving two size variants each and two cases involving three size variants each, **Supplementary Table SI**). We reasoned that CD8^+^ T cell recognition of these identically restricted size variants could either involve cross-recognition of shared epitope structure(s) by the same TcR(s), or involve recognition of genuinely different epitope structures by different TcR(s). To evaluate this, tetramers of the different epitope size variants and the relevant HLA restriction elements were produced with unique fluorochrome labels and used to determine whether the epitope size variants were recognized by the same or different T cell populations. In some cases, distinctly defined and shared subpopulations were observed (**Figure 6 A, D, I, and J**) indicating that one or more unique shared epitope structures were presented and recognized; in other cases, we observed shared subpopulations merging with populations that were single-stained with one of the length-variant tetramers (**Figure 6 C, G, H, I**); and finally, in some cases, no shared subpopulations were observed suggesting that the corresponding length-variants were presented and recognized as being distinctly different **(Figure 6 B, E, F)**. Accommodating length-variants by extending the peptide-binding groove or by one or more aa’s bulging out of the groove (34) could affect the presented epitopes dramatically, whereas accommodating length-variants by protruding out of the N- or C-terminal ends of the groove could leave the non-protruding end of the epitope unaltered. Elucidating the structural basis of these various recognition modes is beyond the scope of this paper.

**Figure 6:**
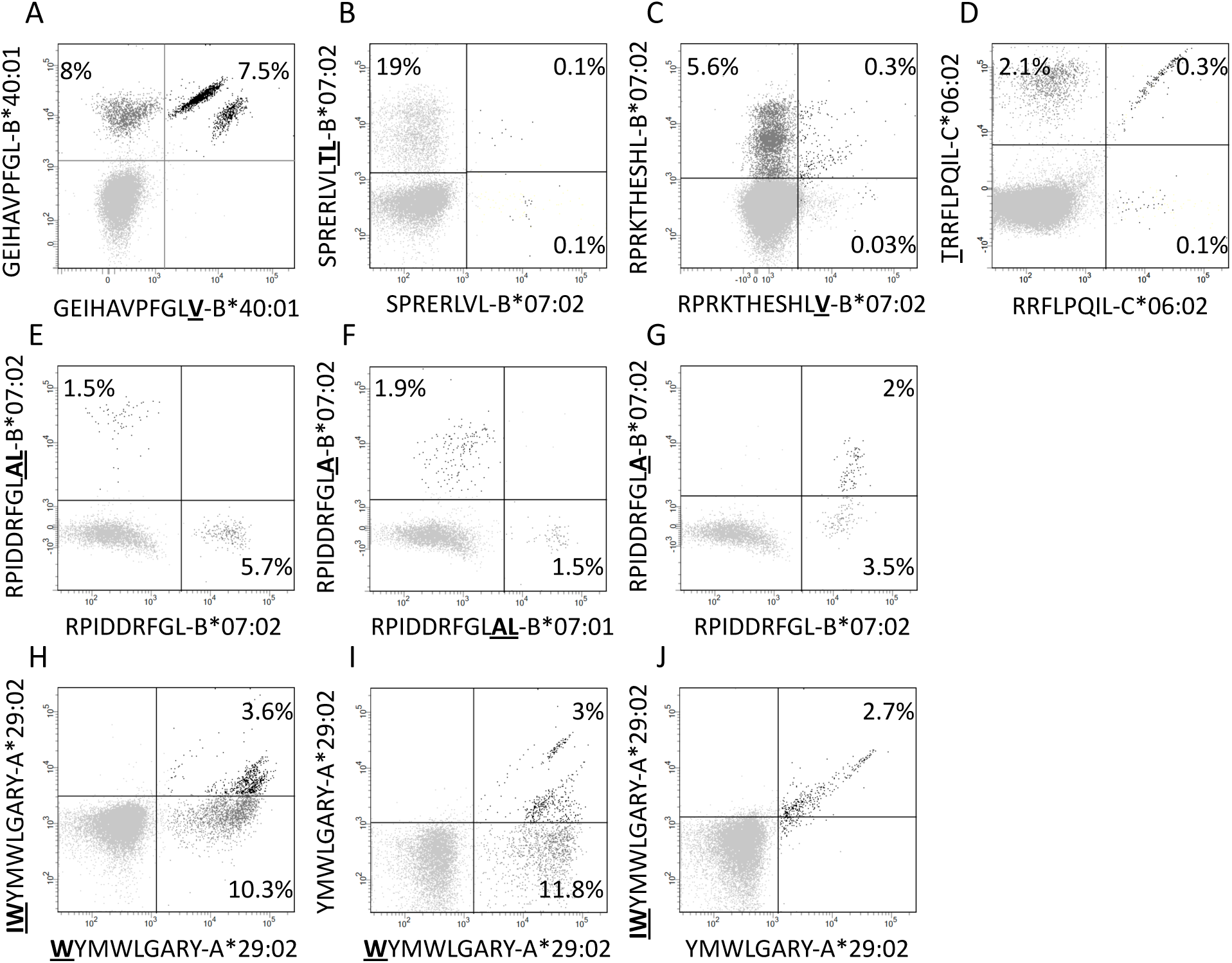
CD8^+^ T cells double stained with combinations of tetramers of size-variants epitopes. PBMCs were *in vitro* expanded for 8 days with relevant 15-mers peptides, and subsequently stained with pairs of tetramers representing size-variants epitopes as indicated in the figure (one member of each pair was PE-labeled and the other was APC-labeled) and analyzed by flow cytometry gating for CD3^+^CD8^+^T cells.

#### 3.5.6. Comparison of strategies of CD8^+^ T cell epitope discovery

One of the more frequent HLA allotypes, HLA-B*07:02, offered an opportunity to compare the HFRI approach with a strictly reverse immunology approach. Theoretically, a total of 13610 peptides of 8-11mer size could be generated from the YFV proteome. NetMHCpan 2.4 predicted 54 of these as being strong HLA-B*07:02 binders at a %Rank of less than 0.5%. We selected 40 of those for further examination (**Supplementary Table SIII**). With one exception, all of these predicted binders supported HLA-B*07:02 tetramer generation, which subsequently were used to examine *ex vivo* obtained PBMC’s from at least 16 HLA-B*07:02^+^ donors. Apart from the epitopes that had already been described (**Supplementary Table SI and SII**), no additional HLA-B*07:02-restricted CD8^+^ T cell epitopes were identified. Thus, a final count can be made: combining the HFRI and a strictly reverse immunology approach, a total of ten unique HLA-B*07:02-restricted CD8^+^ T cell epitopes were found; the HFRI strategy identified nine of these, whereas the reverse immunology strategy identified eight; seven (70%) of these epitopes were shared. Assuming that the number of true positive HLA-B*07:02 epitopes is ten, then both strategies were sensitive (correctly identifying 80-90% of the ten epitopes) and at the same time very specific (correctly rejecting ≈99.6% of the 13600 non-epitopes); the HFRI approach being slightly more sensitive and specific than the reverse immunology approach. The major performance difference between the two strategies arose from the lower false discovery rate (FDR) where the HFRI screening strategy required 13 peptides to identify nine of the ten epitopes found (a FDR of 17%; albeit some of these apparently false positive peptides were eventually identified as epitopes restricted by other HLA-I restricting elements expressed by the donors suggesting that the true false discovery rate of the HFRI approach was even smaller), whereas the reverse immunology approach required 54 peptides to identify eight of the ten epitopes suggesting a false discovery rate of 85%.

In conclusion, the present HFRI approach ranks epitope at the very top of the list of candidates while decimating the false discovery rate (further comparisons of HFRI vs reverse immunology is described in the **Discussion** and detailed in **Supplementary Results and Discussion**).

#### 3.5.7. Efficiencies of CD8^+^ T cell epitope predictors

The unbiased nature of our cohort of 120 different HFRI-identified peptide HLA-I combinations covering 40 HLA-I restriction elements provided an opportunity to evaluate the performance and discriminatory power of various prediction methods such as the authoritative NetMHCpan (both the contemporary version 2.4 (37) and the most recent version 4.0 (38) trained on both eluted ligands (EL) and peptide binding affinity (BA)), and the recent MHCFlurry (39) (trained either only on BA data or on both EL and BA data) (39) and MixMHCpred (trained only on EL data) (40). In addition to these peptide-HLA-I affinity predictors, we also included a stability predictor, NetMHCStabpan 1.0 (41). For each of these methods, predictions scores of all 13610 peptides of length 8-11 aa that could be generated from the 3411 aa YFV proteome were predicted for the relevant HLAs (using %Rank scores allowing comparisons across HLA allotypes and predictors as read-outs). Subsequently, a Receiver Operating Characteristics (ROC) analysis was performed and the Area Under the Curve (AUC) was determined. A non-discriminatory predictor has an AUC of 0.5, whereas a perfectly discriminating predictor has an AUC of 1.0. Applied to this unbiased and validated set of epitopes, all of these predictors gave highly discriminatory AUC’s of 0.98743 to 0.99797 (**Supplementary Figure S2**). These impressive AUC’s are heavily influenced by the many non-immunogenic peptides being correctly rejected; however, this may still leave considerable room for false positive discovery rates (FDR). In this case, a more FDR-averse way to visualize the performance is to use the Frank score, which is the number of false positive predictions (FP) relative to the total number of peptides (N) that can be generated from the source protein (i.e. Frank = FP/N). A Frank score of 0 indicates a “perfect prediction” where a true epitope receives the highest prediction value of all peptides within the source protein and avoids any false positive predictions, whereas a Frank score of 0.5 indicates a random prediction where half of the predictions are false positives. Frank values were calculated for each epitope-HLA pair and predictor (**Supplementary Figure S2**). The best predictors were NetMHCpan 4.0 EL and MixMHCpred, which respectively scored 21 and 20 “perfect” predictions, obtained an average Frank score of 0.001875 and 0.003809, and a median Frank score of 0.000405 and 0.000588, respectively. The median, being a more “outlier-resistant” measure, would respectively indicate that the NetMHCpan 4.0 EL and MixMHCpred methods would place 6 and 8 false-positive non-epitopes ahead of each epitope, corresponding to a false discovery rate of 85 and 89%. These numbers should be appreciated in the context of a random predictor, which would yield a FDR of 99%, and a prefect predictor which would yield an FDR of ≈50% (assuming that only 50% of HLA-presented peptides are immunogenic (2). In line with earlier work (42), comparing the predictive power of the various predictors in terms of the Frank values, NetMHCpan 4.0 EL was found to significantly outperform all other predictors (P < 0.02 in all cases, Wilcoxon matched-pairs signed rank test) (**Supplementary Figure S2**).

### 3.6. CD4^+^ T cell epitope discovery

#### 3.6.1. Identification and validation of CD4^+^ T cell epitopes (exemplified by donor YF1067)

In donor YF1067, the ICS-based screening analysis identified thirty-one 15mer peptides as stimulating CD4^+^ T cell responses. At face value, these 15mer peptide sequences qualified as CD4^+^ T cell epitopes (the IEDB epitope curation manual 2.0 defines a CD4^+^ T cell epitope of 15 residues or less in length as an “exact epitope”). To identify the underlying HLA class II restriction elements, the binding of each of the thirty-one 15mer peptides to each of the HLA-DR molecules of donor YF1067 (*in casu* HLA-DRB1*13:02, -DRB1*15:01, -DRB3*03:01 and -DRB5*01:01) was tested in a biochemical binding affinity assay (43). Nine (28%), eleven (34%), four (13%) and four (13%) of the epitopes bound with an affinity better than 50 nM to one, two, three and four of the donor’s HLA-DR molecules, respectively (**Table II**), whereas three (9%) bound to none of them. Secondly, we generated tetramers for 50 of the (9×1 + 11×2 + 4×3 + 4×4) = 59 strongly interacting peptide-HLA combinations and used these to label *in vitro* expanded CD4^+^ T cells from donor YF1067. Twenty-two of the 50 tetramers successfully identified CD4^+^ T cell epitopes and their HLA-DR-restriction elements (**Figure 7)**. The final validation and enumeration of specific CD4^+^ T cell was performed by an *ex vivo* IFNγ ELISpot analysis (**Table II**). In one case, the same epitope was presented by two different HLA-DRB allotypes and should therefore only be counted as epitope once. Thus, 21 of the 31 different HLA-DR-restricted CD4^+^ T cell epitopes observed in donor YF1067 were identified at the tetramer level; the remaining eleven epitopes were not resolved. The latter could potentially be explained as being restricted by HLA-DQ or DP molecules; something that could not be readily addressed by our tetramer capabilities at the time; albeit, in one case, we successfully generated a NS4B_233-247_-DPA1*01:03-DPB1*04:01 tetramer and identified an HLA-DP-restricted epitope. Although 19 of the 32 CD4^+^ T cell stimulatory peptides bound to more than one of the four HLA-DR allotypes of donor YF1067, there was only one epitope that exploited more than one of the available HLA-DR allotypes as restriction element: the NS5_551-565_ epitope, which was recognized by CD4^+^ T cells in the context of both HLA-DRB1*15:01 and HLA-DRB5*01:01. That this was not a case of TcR cross-recognition was shown by double staining with the two tetramers showing two distinctly different CD4^+^ T cell populations recognizing the NS5_551-565_-epitope presented by either HLA-DRB1*15:01 or HLA-DRB5*01:01 (see section 3.6.3 below). Thus, in donor YF1067, a total of 31 CD4^+^ T cell epitopes were identified; 22 of these could be HLA-DR or -DP tetramer validated.

**Figure 7:**
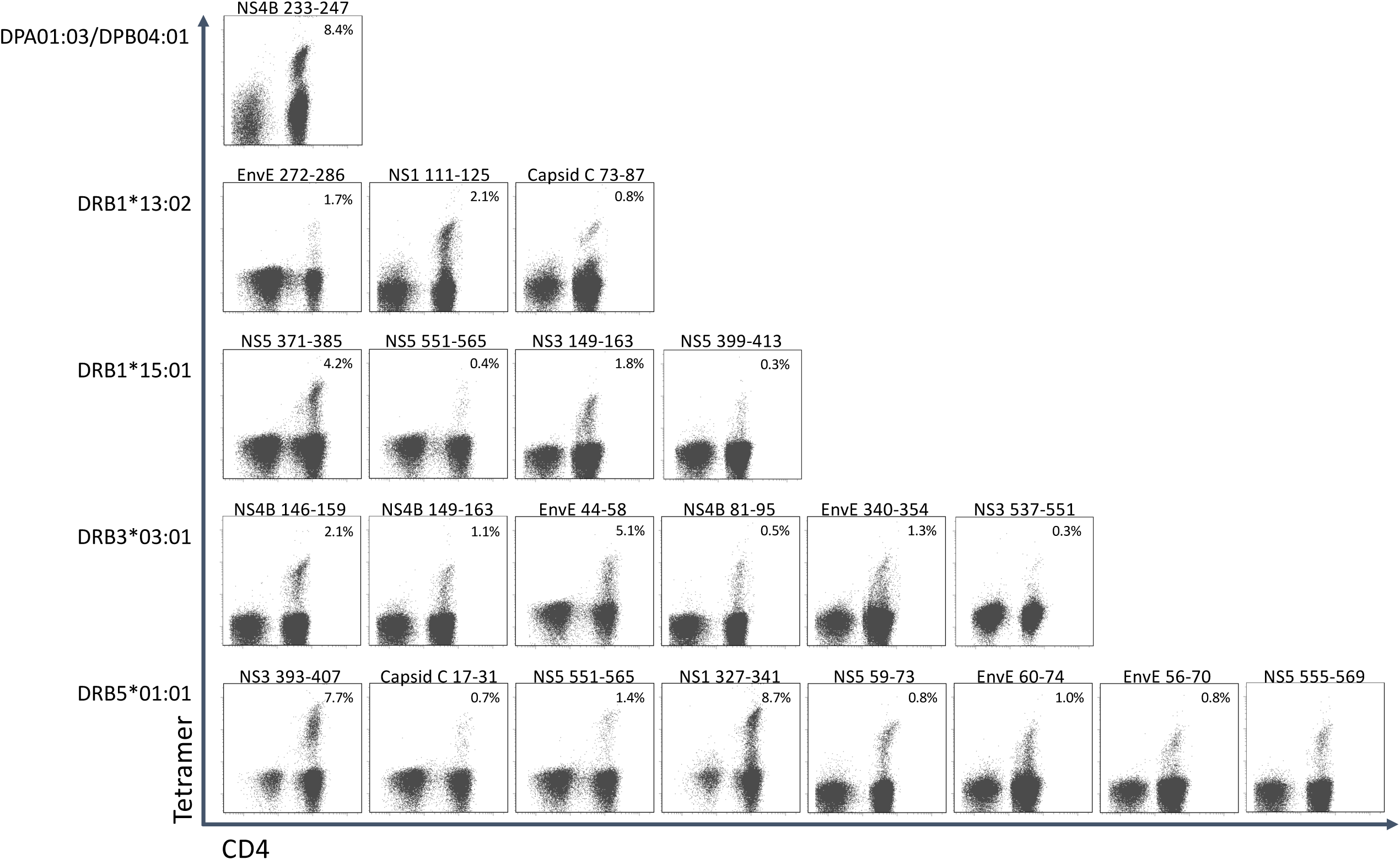
CD4+ T cell epitopes in donor YF1067. PBMC’s from donor YF-1067 were expanded using the stimulatory 15mer peptides identified. For each peptide, a biochemical HLA class II binding assay was used to identify which of donor YF1067’s HLA-II molecules could bind the peptide and therefore could serve as restriction elements. Productively interacting peptide-HLA-II combinations were used to design and generate peptide-HLA class II tetramers. The resulting tetramers were used to stain and analyze expanded CD4^+^ T cells by flow cytometry gating on CD3^+^ CD4^+^ T cells. The identities of the epitopes and their restricting HLA-II elements are indicated.

#### 3.6.2. Extending CD4^+^ T cell epitope discovery to 50 primary YFV vaccinated individuals

The CD4^+^ T cell epitope discovery strategy was extended to the same 50 donors used for CD8^+^ T cell epitope discovery. A total of 192 CD4^+^ T cell stimulatory 15mer epitopes were identified (**Figure 8, Supplementary Table SIV**). Some of these epitopes were frequently recognized. Thus, the single most recognized CD4^+^ T cell epitope, EnvE_44-58_, was recognized in 22 (71%) of 31 tetramer-tested donors tested, and another 12 epitopes were recognized in 10 to 16 (32-52%) of 31 donors. However, most of the 192 epitopes were much less frequently recognized; in fact, 76 of the peptides were recognized in only one (3%) of the 31 donors. We suggest that the strongest and most immunodominant CD4^+^ T cell epitopes have been found.

**Figure 8:**
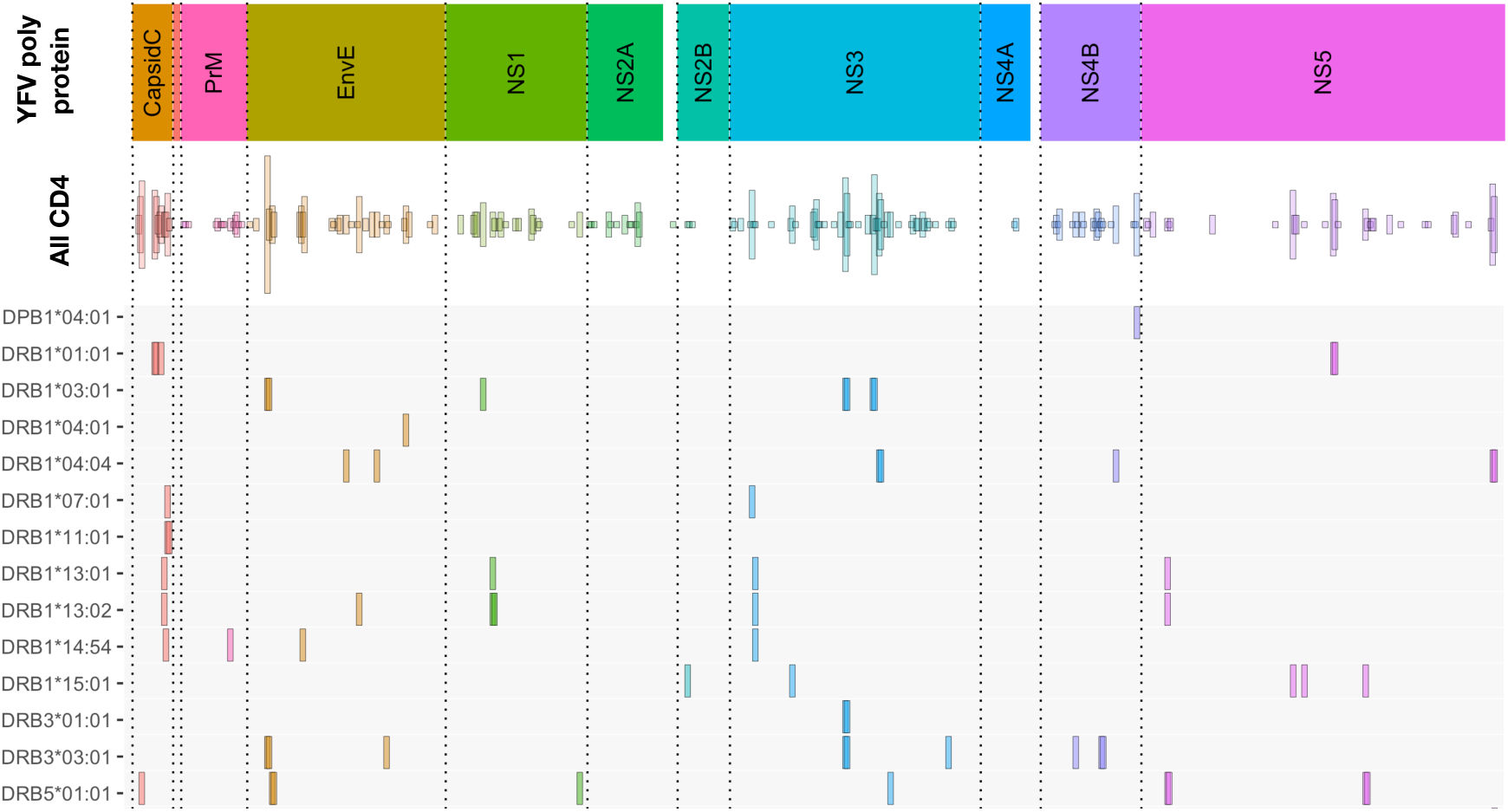
Overview of CD4^+^ T cell epitopes discovered in 50 primary YFV vaccinated donors. Similar to figure 3, the color-coded upper bar indicates the YFV polyprotein. The “all CD4” bar indicates the positions of each of the 192 CD4^+^ T cell stimulatory 15mer peptides and their prevalence. The lower HLA-II allotype-designated bars indicate the tetramer-validated epitopes and their HLA-II restriction elements. The details of each epitope (epitope sequence, CD4^+^ T cell stimulation, ex vivo tetramer staining frequency, and response prevalence is given in **Supplementary Table SV**)

An important objective was to identify and validate the HLA-DR restriction element(s) used to present these epitopes (**see Figure 8 and Supplementary Table SV**). We have evaluated the restriction elements for 74 of the 192 epitopes. For each epitope, the most likely HLA-DR restricting element was selected based on its affinity to one or more of the HLA-DR allotypes available to the donor. Guidance was also obtained from which HLA-DR allotypes were shared amongst the epitope-responding donors. In some cases, more than one strong binding HLA-DR allotype and/or more than one shared HLA-DR allotype were found highlighting that multiple HLA-DR allotypes would have to be considered as potential restriction elements.

In total, 152 peptide-HLA-DR tetramers were generated and used to validate the CD4^+^ T cell epitopes. Of these, 64 tetramers were tested positive for CD4^+^ T cell staining in one or more donors. This covered 50 CD4^+^ T cell peptide epitopes restricted by 13 different HLA-DR molecules (some epitopes were presented by more than one HLA-DR allotype) and one HLA-DP molecule. For 17 of the 50 epitopes, the HLA-DR molecules available to us for tetramer generation did only partially cover the HLA-DR molecules observed in one or more of the responding donors. As an example, the most frequently recognized epitope, EnvE_44-58_, was found in 23 donors (**Supplementary Table SIV and SV**). Using appropriate tetramers, two restriction elements, HLA-DRB1*03:01 and - DRB3*03:01, were identified, however, four of the EnvE_44-58_ responding donors expressed neither the DRB1*03:01 nor the DRB3*03:01. This suggested that one or more additional, not yet identified, restriction element(s) existed for this epitope; something that could apply to more of the 17 epitopes.

#### 3.6.3. Recognizing the same CD4^+^ T cell epitope presented by two to three different HLA-DR allotypes

Some of the 15mer peptides could stimulate CD4^+^ T cell responses restricted by two or three different HLA-DR restriction elements (**Supplementary Table SV**). No donor happened to possess three appropriate HLA-DR molecules, but some did possess two and could generate appropriate CD4^+^ T cell response restricted by both of these restriction elements. In these cases, staining CD4^+^ T cells with two uniquely labeled tetramers, representing either of the two restriction elements, allowed us to address whether the same epitope presented by two different restriction elements were recognized by the same, or by distinctly different, CD4^+^ T cells. When presented by different HLA-DR molecules, five of the eight epitopes (NS5_551-565_ presented by HLA-DRB1*15:01 and - DRB5*01:01 (26 amino acid differences); NS3_285-299_, NS3_281-295_, and EnvE_44-58_ presented by HLA-DRB1*03:01 and -DRB3*03:01 (13 amino acid differences); and Capsid_81-95_ presented by HLA-DRB1*07:01 and -DRB1*11:01 (25 amino acid differences)) engaged distinctly different CD4^+^ T cell populations (**Figure 9A-E**). The remaining three epitopes were presented by the closely related HLA-DR allotypes (HLA-DRB1*13:01 and -DRB1*13:02 (one amino acid difference, a V86G, a part of the peptide binding site interacting with P1 of the core sequence)), NS3_57-71_, NS5_59-73_, and NS1_111-125_, showed various degrees of cross-recognition. The NS3_57-71_ peptide presented by HLA-DRB1*13:01 and -DRB1*13:02 is mostly recognized by separate T cell populations; only a small population recognized the peptide presented by both molecules (**Figure 9F**). For peptides, NS5_59-73_ and NS1_111-125_ about half of the T cells recognizing the peptides presented by HLA-DRB1*13:01 cross-recognized the peptides presented by HLA-DRB1*13:02, with none or a very small T cell population recognizing the peptides presented only by HLA-DRB1*13:02 (**Figure 9G and H**). We speculate that a peptide presented by two restricting HLA-DR molecules with only a few polymorphic amino acid differences may be cross-recognized by some, but not necessarily all, CD4^+^ T cells of appropriate specificity, whereas presentation by two restricting HLA-DR molecules with many polymorphic amino acid differences are more likely to be recognized as being distinctly different.

**Figure 9:**
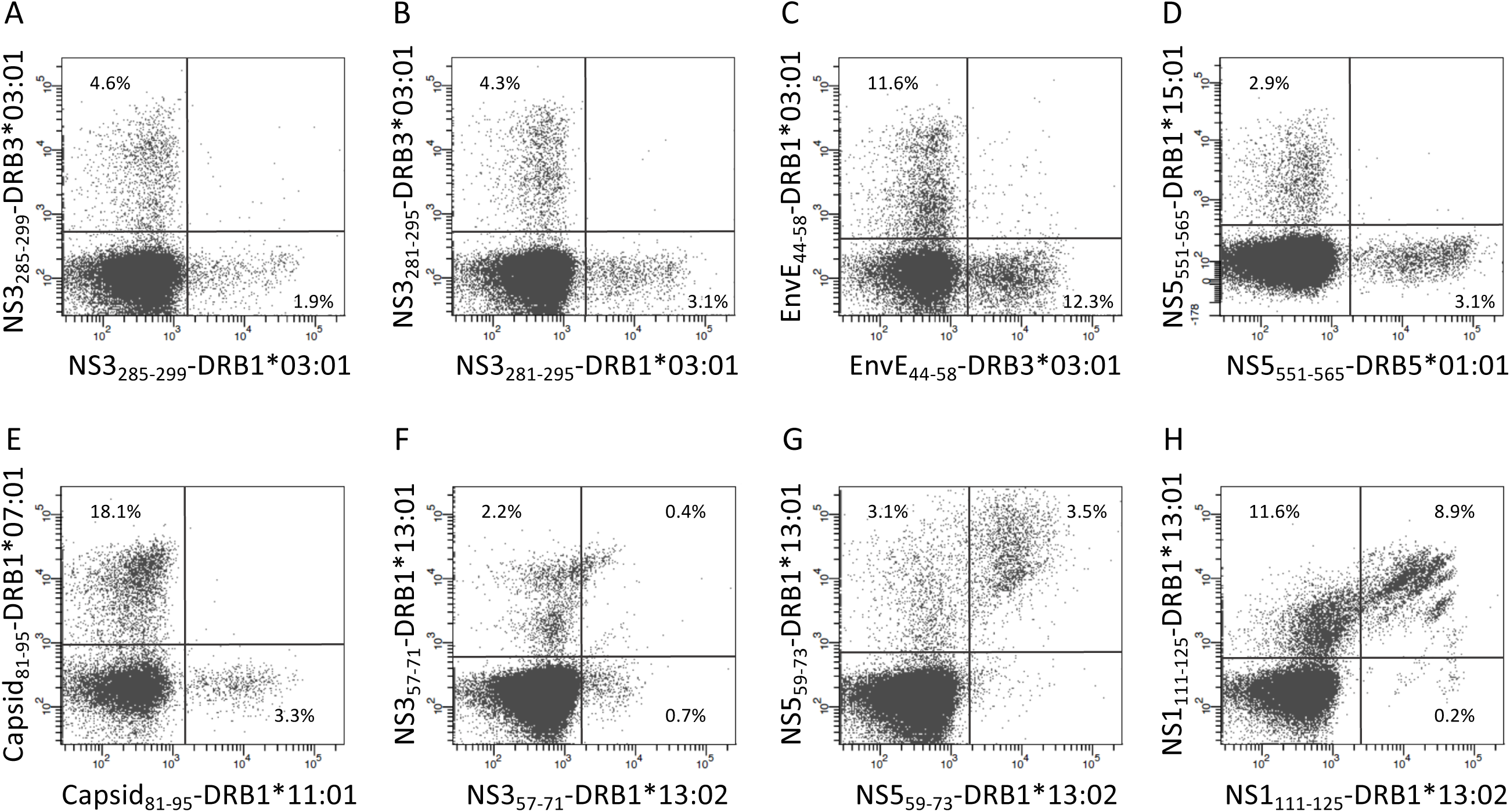
Double staining of CD4^+^ T cells with tetramers representing the same epitope presented by two different HLA-II molecules. PBMC’s from donors recognizing the same epitope presented by two of the HLA-DR molecules of the donors were expanded using the relevant 15mer peptides. The cells were subsequently double stained with the indicated two tetramers representing peptide-HLA-DR combinations labeled with PE or APC, respectively, and analyzed by flow cytometry gating on CD3^+^CD4^+^ T cells. The PE-labeled tetramers are shown on the x-axis and the APC-labeled tetramers are shown on the y-axis.

#### 3.6.4. The recognition of overlapping CD4^+^ T cell epitopes presented by the same HLA-DR allotype expands the diversity of CD4^+^ T cell specificities

In 16 cases, two consecutive overlapping 15mer peptides stimulated CD4^+^ T cell responses restricted by the same HLA-DR restriction element. If the two peptides of such an overlapping 15mer peptide pair were presented through two different core regions, one for each, then the two neighboring epitopes should be perceived as being distinctly different and should be recognized by two disparate CD4^+^ T cell populations. Alternatively, if the two peptides were presented through the exact same core region, then the two neighboring epitopes could potentially be perceived as being identical and be recognized by the same CD4^+^ T cell populations. To examine this, CD4^+^ T cells were double-stained with HLA-DR tetramers, which had been prepared with each of the overlapping peptides of a 15mer pair and labeled with a unique fluorochrome. We analyzed 10 such pairs and found a wide variety of staining patterns. In no case did two peptides of an overlapping pair engage two distinctly different CD4^+^ T cell populations; rather, in all cases observed, the two peptides engaged at least some shared CD4^+^ T cell populations suggesting usage of shared core regions. In most cases, a plethora of shared, yet subtly different, CD4^+^ T cell populations were observed (**Figure 10**). By way of examples, tetramers representing the overlapping HLA-DRB1*01:01-restricted 15mer peptides, CapC_49-63_ and CapC_53-67_, revealed multiple distinct CD4^+^ T cell subpopulations, which recognized one, the other, or both tetramers at various efficiencies (**Figure 10A**); whereas tetramers representing the overlapping HLA-DRB1*01:01-restricted 15mer peptides, NS5_471-485_ and NS5_475-489_, revealed almost exclusively CD4^+^ T cell subpopulations recognizing both tetramers, albeit clearly comprising multiple distinct subpopulations (**Figure 10B**). We argue that this phenomenon increases and diversifies CD4^+^ T cell responses.

**Figure 10:**
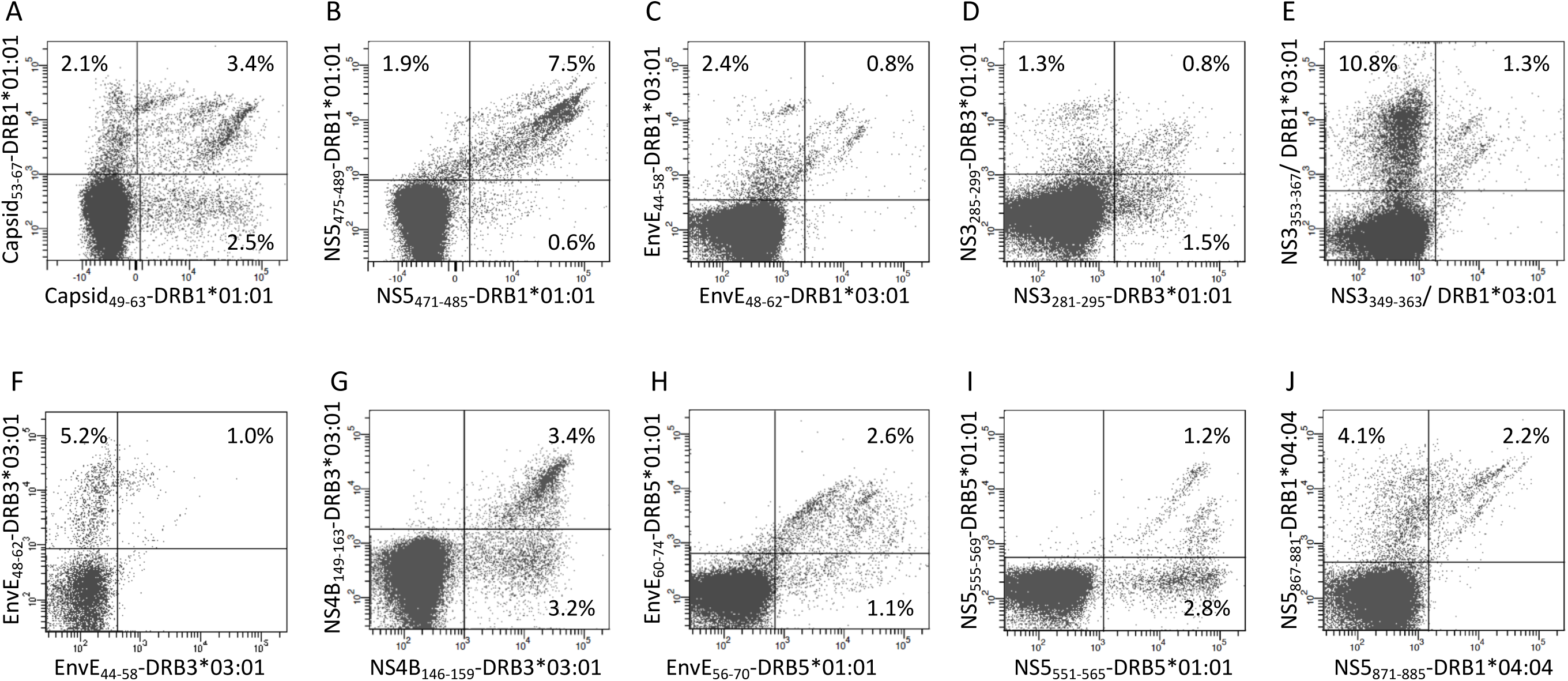
Double staining of CD4^+^ T cells with tetramers representing two overlapping 15-mer epitopes presented by the same HLA-II molecule. PBMC’s from donors recognizing two overlapping 15mer epitopes presented by the same HLA-DR molecule were expanded using the relevant 15mer peptides. The cells were subsequently double stained with the indicated tetramers representing two overlapping peptide-HLA-DR combinations labeled with PE and APC, respectively, and analyzed by flow cytometry gating on CD3^+^CD4^+^ T cells. The PE-labeled tetramers are shown on the x-axis and the APC-labeled tetramers are shown on the y-axis.

### 3.7. Distribution of YFV-specific CD8^+^ and CD4^+^ T cell epitopes

Apart from two small proteins, the 20 aa ER anchor and the 164 aa prM proteins, all YFV proteins contained both CD8^+^ and CD4^+^ T cell epitopes. On average, the frequencies of CD8^+^ and CD4^+^ T cell epitopes were ≈3 and 6 per 100 aa, respectively (**Table III**). Notably, the CD8 T cell epitopes, which have been tetramer mapped exhaustively, exhibited stretches of overlapping epitopes restricted by several different HLA class I molecules: twelve stretches encompassing two epitopes, five stretches encompassing three epitopes, and three larger hot-spots areas encompassing four to six epitopes, many of which were presented by several different HLA molecules. Thus, the frequently recognized EnvE_200-240_ sequence comprised six peptide epitopes and ten HLA-restriction elements giving a total of twelve epitope-HLA combinations (**Figure 11**). These three hot-spot regions accounted for about 15 of the 97 (15%) CD8^+^ T cell epitopes identified, and encompassed 15 HLA-I restriction elements covering ≈ 77% of the Caucasian race. Although the YF protein was generated as one long precursor polyprotein, no epitopes were found in any of the overlaps between the different processed proteins.

**Yable III:** Overview of the number of the total number of YFV-specific CD4^+^ and CD8^+^ T cell epitopes discovered. The frequencies of epitopes per 100 aa are also given.

| **YFV proteins** | **Protein length**<br>**(amino acids)** | **CD4<sup>+</sup> T cell epitopes**<br>**Number**<br>**(frequency)** | **CD8<sup>+</sup> T cell epitopes**<br>**Number**<br>**(frequency)** | **Total CD4+ and CD8+**<br>**T cell epitopes**<br>**Number (frequency)** |
| --- | --- | --- | --- | --- |
| **Capsid C** | 101 | 12 (11.9) | 2 (2.0) | 14(13.9) |
| **prM** | 164 | 14 (8.5) | 0 (0.0) | 14 (8.5) |
| **Envelope** | 493 | 29 (5.9) | 15 (3.0) | 44 (8.9) |
| **ER anchor** | 20 | 0 (0) | 2 (10.0) | 2 (10.0) |
| **NS1** | 352 | 18 (5.1) | 12 (3.4) | 30 (8.5) |
| **NS2A** | 224 | 13 (5.8) | 8 (3.6) | 21 (9.4) |
| **NS2B** | 130 | 4 (3.1) | 3 (2.3) | 7 (5.4) |
| **NS3** | 623 | 47 (7.5) | 19 (3.0) | 66 (10.6) |
| **NS4A** | 149 | 2 (1.3) | 6 (4.0) | 8 (5.4) |
| **NS4B** | 250 | 21 (8.4) | 5 (2.0) | 26 (10.4) |
| **NS5** | 905 | 31 (3.4) | 25 (2.8) | 56 (6.2) |
| **Total YF polypeptide** | 3411 | 191 (5.6) | 97 (2.8) | 288 (8.4) |

**Figure 11:**
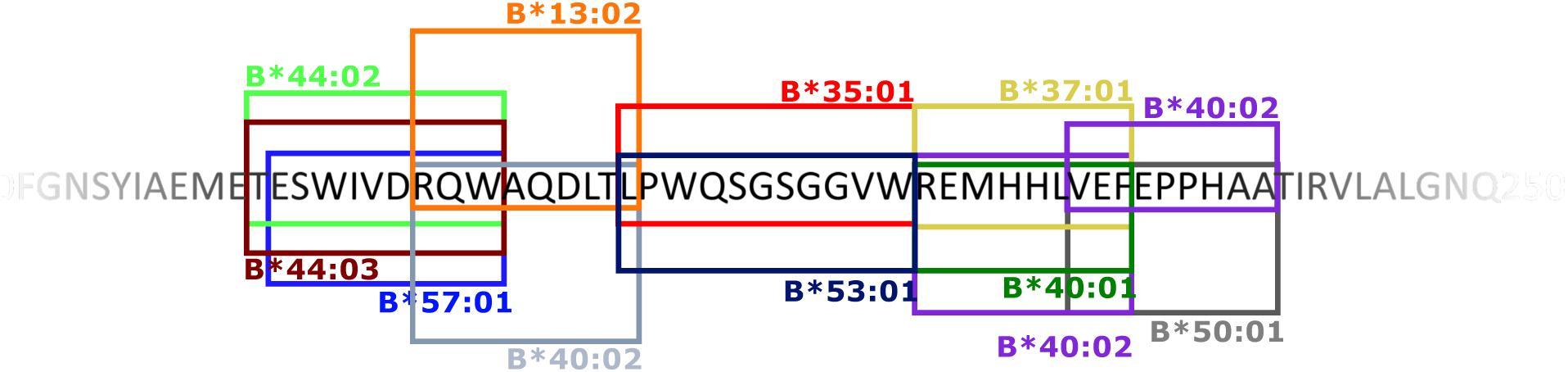
An example of a CD8^+^ T cell epitope hot spot: the EnvE_200-240_ region. The 40 amino acid sequence EnvE 200-240 included 6 different CD8+ T cell epitopes presented by 10 different HLA-I molecules – in total 12 different peptide-HLA-I combinations. The boxed sequences represent the epitopes that are presented and recognized by the restriction elements indicated at the top or bottom of the box. A box and its indicated restriction element is identically colored.

## 4. Discussion

In order to understand the complexity of the human T-cell response to a circulating pathogen, and its potential impact on population dynamics of both pathogen and host, knowing a wide range of epitopes relevant for T-cell/pathogen interplay is essential. However, identifying the exact epitope sequence and the exact HLA allotype involved in T cell recognition of a specific pathogen is a demanding challenge. Over the years, a plethora of methods have been used to identify T cell epitopes. There are two major and principally different approaches of T cell epitope discovery. The “forward immunology” approach (28, 29) uses specific T cell responses as a starting point to search for the underlying T cell epitope and its MHC restriction, whereas the “reverse immunology” approach (30, 31) uses predictions (e.g. of peptide-MHC interactions) to suggest possible T cell epitopes and then screen them for their ability to stimulate specific T cell responses (reviewed in 9, 44, 45). The experimental procedures involved in both of these epitope discovery modes tend to involve slow, low throughput, cumbersome and expensive processes (e.g. expression cloning of antigen libraries and/or HLA genes (28, 46-49)), synthesis of peptide libraries etc.). In contrast, the bioinformatics component of a reverse immunology approach offers a process that is fast, of high capacity and throughput, yet very easy and inexpensive; a process, which is well-suited to support systematic analyses of genomic and proteomic information (3, 4, 9, 30). It is not surprising that reverse immunology has become the preferred approach to T cell discovery. The need for high speed and capacity is of obvious importance in emerging infectious diseases (including bioterrorism), and even more so in personalized cancer immunotherapy where fast and high-throughput methods are essential for the selection of relevant and safe cancer neoepitopes in real time. Current peptide-MHC predictors are highly sensitive and specific (96.5% and 98.5%, respectively (16)). However, despite continued improvements of these predictors, the false discovery rate (FDR) is very high (8, 10, 18); something that compromises the successful inclusion of one, or preferably more, T cell epitopes in cancer immunotherapy even if these encompass up to 10-20 predicted epitopes (20, 21). Reducing the FDR while maintaining the sensitivity will be needed if reverse immunology in the future should fully support neoantigen discovery and secure timely, personalized immunotherapy of cancer (19). Indeed, most of the larger CD8^+^ and CD4^+^ T cell epitope submissions to the IEDB have been identified by “reverse immunology”. Thus, Sette and coworkers used “reverse immunology” to identify Dengue virus-specific T cell epitopes and have, as of July 2019, contributed with the single largest submissions of CD8^+^ and CD4^+^ T cell epitopes (IEDB reference ID 1027503, 1031475 and 1031301). In contrast, the “forward immunology” approach has fallen relatively into disuse. An innovative approach pioneered by Koelle and co-workers, which has resulted in larger IEDB submissions of CD8^+^ and CD4^+^ T cell epitopes (e.g. IEDB reference ID 1021375), have used a “forward” component where co-transfecting panels of APC with cDNA encoding antigen and HLA class I or II, each APC representing a single antigen and a single HLA restriction element, were used to interrogate CD4^+^ and CD4^+^ T cell responses of virus infected donors (49). The “forward” component of this approach identified intact immunogenic protein antigens and their restriction element(s); however, for the epitope discovery part of this work, the entire antigen was subjected to a “reverse” component predicting the epitope(s) and its HLA restriction element(s). Another innovative approach, Tetramer Guided Epitope Mapping (TGEM), pioneered by Kwok and James, which has resulted in large CD4^+^ T cell epitope submissions (IEDB references ID 1026930 (50), 1013360, 1016040, and 1020783), have also used a “forward” component. Longer overlapping peptides representing entire antigens were offered to single HLA class II molecules and the resulting peptide-HLA class II complexes were multimerized and the ensuing tetramers used to interrogate CD4^+^ T cell responses of appropriate donors. Using shorter overlapping peptides suitable as CD8^+^ T cell epitopes, Maeurer and coworkers established a tetramer-based approach for CD8^+^ T cell epitope discovery, which also resulted in larger IEDB submissions (IEDB ID 1026840 (51)). This latter approach would obviously be very peptide intensive if every relevant peptide was to be tested in that way (e.g. the Yellow Fever proteome would require 13610 peptides to represent all possible 8-11mer peptides). Here, we have generated a “hybrid forward-reverse immunology” (HFRI) approach capable of doing concurrent CD8^+^ and CD4^+^ T cell epitope discovery and demonstrated that it can perform large-scale epitope discovery and at the same time decimate the false positive discovery rate. For the initial “forward immunology” screen, we used an overlapping peptide library of 850 15mer peptides overlapping by 11 aa, which represented the entire 3411 aa Yellow Fever Virus proteome, to stimulate PBMC’s obtained *ex vivo* from primary Yellow Fever Virus vaccinees at the peak of the resulting T cell response. In itself, this overlapping 15mer peptide library represented all possible YFV-specific, CD4^+^ T cell epitopes of up to 12 aa in length. In addition, since 15mer peptides are further processed during *in vitro* ICS and/or ELIspot assays, it also represented all shorter YFV-specific, CD8^+^ T cell epitopes (a total of 13610 peptides of length 8-11 aa, which can be generated from the YFV proteome). Distributing this peptide library into 4 matrices, the initial screening effort could be reduced to testing ≈120 peptide pools for their ability to stimulate both CD8^+^ and CD4^+^ T cell responses. The matrix design subsequently allowed us to home in on the individual T cell stimulatory peptides. The subsequent “reverse immunology” approach was applied to all 15mer peptides containing CD8^+^ T cell epitopes. In silico, the affinities of all possible 8-11mer peptides that could be generated from the 15mer were predicted in the context of up to 6 different HLA-A, -B and -C allotypes per individual. This reduced the number of potential peptide-HLA-A, -B or -C combinations from 156 per stimulatory 15mer peptide to typically one to three combinations. The most likely peptide binders were synthesized and used to generate appropriate peptide-HLA-I tetramer(s), which subsequently were used to validate CD8^+^ T cell epitope(s). For the vast majority of T cell stimulatory 15mer peptides, at least one epitope was identified per 15mer peptide. Once the stimulatory 15mer peptides had been identified, predicting the exact epitope and its restriction element was a highly efficient process; typically, the epitopes ranked first, second or third amongst the many potential epitope-HLA combinations. As a cost-saving measure, if the predictions clearly discriminated between the candidates, a consecutive process was applied whereby the top peptide(s) were synthesized and tested before any next tier peptides were synthesized and tested. This HFRI approach was extended to 50 primary YFV vaccinees, where it identified and tetramer-validated 92 CD8^+^ T cell epitopes (predominantly of size 9 to 10mer, range 8 to 11mer) covering 40 HLA-I allotypes (representing a total of 120 peptide-HLA-I combinations). Before this work, the IEDB had registered ten YFV-specific CD8^+^ T cell epitopes as being “exact epitopes” (i.e. length from 7 to 11 aa) and restricted by an HLA allotype defined at high (4-digit) resolution; however, none of them were tetramer validated. Four of the ten already registered YFV-specific, CD8^+^ T cell epitopes were included in the 92 epitopes identified here. Thus, the present approach identified and validated 92 – 4 = 88 new, or ≈90% of all currently known, YFV-specific, CD8^+^ T cell epitopes. The total number of exact CD8^+^ T cell epitopes with high resolution HLA-I restriction, which are currently registered in the IEDB is 2612 of which 1101 have been tetramer validated (extracted from the IEDB, July 2019). Thus, this study accounts for >8% of these tetramer-validated human CD8^+^ T cell epitopes.

To evaluate the prevalence of the different YFV-specific CD8^+^ T cell immune responses, the tetramer analysis was extended to additional vaccinees with the appropriate HLA-I allotypes. Many epitopes were frequently observed (i.e. were highly prevalent) in vaccinees with the appropriate HLA allotype. Thus; 25 (≈31%) and 50 (≈62%) of 81 CD8^+^ T cell epitopes were observed in ≥90% and ≥50%, respectively, of vaccinees with the appropriate HLA-I allele. Conversely, 18 (≈75%) of 24 HLA-I allotypes presented at least one CD8^+^ T cell epitope with a prevalence of ≥90%. Thus, the HFRI approach identified a cohort of immunodominant Yellow Fever-derived peptides, which could be of broad diagnostic and therapeutic interest. Large-scale T cell epitope discovery could also address more fundamental issues in immunobiology. Pertinent examples of phenomena that are poorly understood include the closely related immunodominance (that the immune response is focused on just a few of the many available determinants expressed by a pathogen) and immunodomination (that the immune response of one specificity can suppress the response of another specificity). Not surprising, these phenomena are closely related to antigen processing and presentation including MHC and T cell repertoire (35). The vast majority of experimental data on immunodominance and immunodomination emanates from studies involving inbreed mice. Few studies in humans address immunodominance (e.g. 52); to the best of our knowledge none involve immunodomination. The latter is particularly difficult to address in an outbreed system like the human where the extremely diverse HLA creates context dependent effects that confounds attempts to address immunodomination. Assuming that the context-dependent effects HLA could even out in larger donor cohorts, we exploited the size of our study to ask whether the presence of HLA-A*02:01, which restricts a strongly immunodominant, NS4B_214-222_-specific T cell response, would correlate with a reduction of responses restricted by other HLA allotypes. Indeed, under these conditions, we could demonstrate such a correlation in the presence of HLA-A*02:01, but not in the presence of HLA-A*01:01 or -A*03:01. Note that the HLA-A*01:01 or -A*03:01 allotypes themselves featured a hierarchy of immunodominant T cell responses i.e. they are valid HLA restricting elements. This may be the first demonstration of primary anti-virus responses being subjected to immunodomination in humans. A further analysis of the mechanism of behind these phenomena is beyond the scope of this paper.

HLA-C restricted, CD8^+^ T cell epitopes were scarcely represented (<3%) in the IEDB; something that potentially could be explained by HLA-C being insufficiently investigated. A priori, we expected that the unbiased nature of our approach would reveal several HLA-C restricted CD8^+^ T cell epitopes, however, we only found one case of a strong and highly prevalent CD8^+^ T cell response, which could not be explained by any of the HLA-A or -B allotypes available to the responding donors. Instead, a strongly predicted binding to a shared HLA-C allotype amongst the responding donors suggested an HLA-C*06:02 restricted response. Eventually, two HLA-C*06:02-restricted epitope length variants; NS3_207-213_ (TRRFLPQIL) and NS3_208-213_ (RRFLPQIL), were tetramer validated. These were the only HLA-C restricted CD8^+^ T cell epitope identified; all other identified CD8^+^ T cell epitopes were validated as being either HLA-A or -B restricted. HLA-C is less polymorphic and is known to be expressed at a lower level than HLA-A and -B (53-56); something that has been correlated with reduced cytotoxic T lymphocyte responses (57, 58). In the case of the HLA-C*06:02-restricted NS3_207-213_ (TRRFLPQIL) epitope identified here, any reduced expression level of HLA-C*06:02 might have been compensated by the very strong predicted binding affinity for NS3_207-213_. Although weaker HLA-C-restricted CD8^+^ T cell responses may have been missed, we would argue that it is unlikely that we have missed strong and prevalent HLA-C restricted CD8^+^ T cell epitopes. Thus, we suggest that the paucity of strong HLA-C restricted CD8^+^ T cell responses, at least in an acute viral infection like yellow fever virus, is not due to HLA-C having been neglected in the scientific literature, but rather reflects a true biological phenomenon. Notwithstanding, future CD8^+^ T cell discovery efforts should include HLA-C, in particular if one or more HLA-C restricted epitopes can be suggested in a situation where there are no obvious HLA-A or -B restricted candidates.

Concurrent with CD8^+^ T cell discovery, the “forward-reverse immunology” approach also allowed HLA-II-restricted CD4^+^ T cell epitope discovery. The initial matrix-driven “forward” analysis of 50 donors identified 192 CD4^+^ T cell stimulatory YFV-derived 15mer peptides. This suggests that CD4^+^ T cell epitopes are more numerous than CD8^+^ T cell epitopes, perhaps as much as 2-3 times greater. If generalizable, this would have important implications for CD4^+^ T cell immunity since, everything else being equal, it would be more difficult for a microorganism to escape many CD4^+^ T cell epitopes than fewer CD8^+^ T cell epitopes. Addressing the number of immunogenic open reading frames, other have also hinted at a greater preponderance of CD4^+^ than CD8^+^ T cell epitopes (59, 60); to the best of our knowledge, ours is the first proteome-wide study that have made this observation at the epitope level.

The identification of the restricting HLA class II element(s) is a serious challenge in part due to different HLA-II allotypes having overlapping peptide binding repertoires (61). In fact, this problem is so manifest that Sette et al have developed a panel of 46 different single HLA-II transfected cell lines to identify HLA-II restriction elements (62). It would be ideal if HLA-II restrictions could be identified by predictors and then validated by tetramer analysis. Unfortunately, the contemporary CD4^+^ T cell epitope discovery tools were immature (e.g. the early NetMHCIIpan predictors were relatively inefficient and focused solely on the HLA-DR isotypes), and access to peptide-MHC class II tetramers was very limited. Moreover, *ex vivo* frequencies of tetramer-positive CD4^+^ T cells tend to be less than 0.01%, which make them difficult to detect. Thus, our CD4^+^ T cell epitope discovery process was not exhaustive; however, as CD4^+^ T cell discovery tools mature, we believe that the efficiency of CD4^+^ T cell epitope discovery eventually should approach that of CD8^+^ T cell epitope discovery.

Here, using a panel of recombinant HLA-DR molecules, we measured the binding affinity of the overlapping 15mer peptides to the most common HLA-DR allotypes. For each stimulatory 15mer peptide, this suggested which of the donor’s HLA-DR molecules should be used to generate peptide-HLA-DR tetramers for validation of CD4^+^ T cell epitopes. This “brute force” approach was extended to 31 donors, where we tetramer-validated 50 CD4^+^ T cell epitopes covering 13 different HLA-DR allotypes (and one HLA-DP allotype). As of July 2019, the IEDB has registered a total of 1915 YFV-specific CD4^+^ T cell epitopes as being “exact CD4^+^ T cell epitopes” (i.e. length 15 aa, or less) and restricted by an HLA-II defined at high (i.e. 4-digit) resolution; 368 of which have been tetramer-validated. Thus, the tetramer-validated YFV-specific CD4^+^ T cell epitopes reported here represents a significant increase in the number of tetramer-validated CD4^+^ T cell epitopes. It should be noted that James and coworkers have identified and tetramer-validated 94 different YFV-specific CD4^+^ T cell epitopes (IEDB reference ID 1026930 (50)) that are 17 aa long and therefore fall just outside the definition of an exact CD4^+^ T cell epitope.

A detailed examination of CD4^+^ T cell responses revealed a phenomenon that could have profound biological and practical implications for CD4^+^ T cell recognition. In many cases, two consecutive overlapping 15mer peptides stimulated CD4^+^ T cell responses, which were restricted by the same HLA-DR restriction element. When the responding CD4^+^ T cells were double-stained with HLA-DR tetramers, which had been prepared with each of the overlapping peptides of a 15mer pair and labeled with a unique fluorochrome, we observed a plethora of different, yet partially shared, CD4^+^ T cell specificities. Situations where overlapping peptides are presented must occur regularly *in vivo* since experiments sequencing natural peptides eluted of HLA-II molecules frequently find large series of staggered peptides surrounding each core region (63). Exploiting this wealth of closely related peptides to engage a large number of different CD4^+^ T cell specificities recognizing the same core region in slightly different ways (something that actually was noted years ago (64)), may represent a biologically significant diversification mechanism of CD4^+^ T cell responses reducing the risk of pathogen escape and increasing the chances of recognizing a given target. This phenomenon is also important for how CD4^+^ T cell responses should be analyzed. A single peptide-HLA-II tetramer is likely to engage a range of T cells of various avidities for the tetramer; something that might explain why single HLA-II tetramers often appear to label a poorly defined, non-base line separated, mono-specific CD4^+^ T cell population of low frequency. If examined with two “overlapping” tetramer, this could in reality turn out to be a heterogeneous collection of better defined and separated CD4^+^ T cell populations of a higher accumulated frequency. From a practical perspective, this implies that double staining involving two overlapping peptide-HLA-II tetramers will be needed to faithfully enumerate and monitor any CD4^+^ T cell response. From a technical perspective, this will increase the observed frequencies of specific CD4^+^ T cells for a given specificity (i.e. increase the sensitivity of the analysis), and it will increase the resolution of the flow cytometric analysis since it may separate various positively staining subpopulations and provide a better discrimination against negatively staining CD4^+^ T cells.

The T cell epitope discovery approach described here has several advantages. In the forward immunology component, an overlapping peptide library is used to search for peptides containing CD4^+^ or CD8^+^ T cell epitopes. In the subsequent reverse immunology component, pan-specific predictors are used to identify the underlying epitope and its HLA restriction element. These steps can be done in any (obviously outbred) individual of any HLA haplotype using simple and standardized conditions. This reduces the number and combinations of peptides and HLA allotypes that should be considered for peptide-HLA tetramer generation and used in the final validation of the discovered T cell epitopes. As shown here, this approach is efficient and, not surprisingly, it reduces the false discovery rate. As peptide-HLA class I and II predictors and tetramer technologies mature, this approach will eventually be able to cover all frequently found HLA class I and II iso- and allotypes. This approach is systematic, all-inclusive, complete, and global in the sense that it includes all protein antigens and peptide epitopes, encompasses both CD4^+^ and CD8^+^ T cell epitopes, and can be extended to all individuals of all populations. This approach could be extended to other attenuated live virus vaccines (e.g. those targeting measles, mumps, rubella, polio, and HPV). Compared to a strictly reverse approach, significant disadvantages of the HFRI approach include the time and cost associated with establishing a complete overlapping peptide library as well as using a cellular readout as an initial selection step. Therefore, this will probably not be justified if the aim is to identify epitopes in an urgent effort involving one donor (e.g. for cancer immunotherapy purposes); rather, it would be appropriate if the aim is to examine a large panel of donors in order to get population-wide data including immunodominance, candidates for diagnostics and vaccine development for infectious disease purposes (examples include a range of emerging and re-emerging infectious diseases like HIV, SARS, MERS, Chikungunya, Dengue, West Nile, Zika, Ebola and SARS-CoV-2). For all of the above examples, the proteomes are small enough that their entire proteomes could be addressed by an overlapping peptide strategy using the number of PBMC’s that could be obtained from a donor. Addressing larger pathogen proteomes (e.g. herpes virus, bacteria or parasites; or smaller highly variable virus like HIV and Dengue) in their entirety would require either a selection process down-sizing the source target protein antigens, or the development of novel miniaturized, yet high-capacity, technologies. One could envision that future investigations of emerging diseases would include population-wide T cell epitope discovery efforts using blood samples from patients, convalescents and/or long-term survivors, which all possess important information on T cell epitopes and responses. Similarly, one could envision that approval and registration of new vaccines could include population-wide analysis of T cell epitopes and responses.

Another important advantage of the forward-reverse approach presented here is the unbiased nature of the T cell epitope discovery process. Whereas current data-driven bioinformatics peptide-MHC predictors are quite accurate, the need for even better predictors stresses not only the need for high-quality training data, but also the need for high-quality validation data. In this context, there is an inherent problem in most epitope discovery efforts being dependent on peptide-MHC predictors since this effectively means that current T cell epitope discovery submissions tend to be biased by current predictors; something that might compromise the validation and benchmarking of predictors. Having reasoned that our forward-reverse approach captures about 90% of the true T cell epitopes, we would like to propose that the resulting data is largely unbiased and should serve as an appropriate benchmark (others have reached similar conclusions (48)). As an example, we used the CD8^+^ T cell epitopes identified here to benchmark current predictors. All current predictors were quite efficient and accurate. The newer predictors, some of which included immunopeptidomics and therefore may also reflect antigen processing, were better than the older predictors (as also noted by others (42)). However, these improvements are incremental and even the newest predictors were afflicted by high FDR’s. Taken together, this could be interpreted as a need for a change in how we predict T cell epitopes that is more fundamental than merely acquiring more peptide-MHC affinity and/or stability data e.g. by including T cell receptor specificities and repertoire propensities. A source of unbiased T cell epitope data would be instrumental in improving predicting tools.

In conclusion, for smaller proteomes, it is possible to design a limited set of overlapping peptides spanning the entire proteome and use these to reveal the vast majority, if not all, specific CD4^+^ and CD8^+^ T cell responses concurrently; use predictors to identify the underlying combination of peptide epitopes and MHC restriction elements; and finally use this information to construct suitable peptide-MHC multimers and validate the T cell epitopes discovered. Performing this in cohorts of patients or vaccinees allows for a systematic, global and cost-efficient analysis of CD4^+^ and CD8^+^ T cell epitopes, and evaluation of their immunodominance.

## 5. Materials and Methods

### 5.1. Study approvals, donors, and YFV vaccination

As previously described (26), this study was approved by the Danish National Committee on Health Research Ethics (protocol # H-1-2009-095), and the collection of data and cells was approved by The Danish Data Protection Agency (permission 2008-41-2732). All volunteers gave written informed consent prior to participation. Based on previous YFV vaccination history and their International Card of Vaccination, healthy volunteers, who for traveling purposes were about to receive a primary YFV vaccination, were recruited. The attenuated YFV vaccine, 17D-204 (Sanofi Pasteur; marketed as Stamaril in more than 70 countries globally and as YF-VAX in the USA) was administered intramuscularly. About 42% of the volunteers received an YFV vaccination only, whereas the remaining 58% received additional vaccines, typically killed, inactivated or subunit vaccines; in no case was the YFV vaccine co-administered with another live attenuated vaccine.

### 5.2. Blood samples and PBMC preparation

As previously described (26), blood samples were obtained just prior to and after the YFV vaccination (typically day 10-20 post vaccination, range 9 to 41 days). Peripheral blood mononuclear cells (PBMC) were isolated by density gradient centrifugation using Ficoll-Paque™ Plus (GE Healthcare Europe, Brøndby, Denmark). They were either examined directly *ex vivo* or cryopreserved in 10% DMSO and 90% FCS at -150°C for later *in vitro* analysis.

### 5.3. High-resolution HLA-typing

Chromosomal DNA was prepared from the PBMC’s and sequence-based typing (SBT) was used to perform high-resolution (i.e. 4 digit) HLA-typing (Genome Diagnostics, Utrecht, The Netherlands). All loci encoding classical HLA molecules were typed i.e. the three class I Ioci, HLA-A, -B, -C and the six class II loci, HLA-DRB1, -DRB3/4/5, -DQA1, -DQB1, -DPA1 and -DPB1.

### 5.4. T cell marker analysis

The PBMCs were analyzed *ex vivo* for the T cell markers, CD3, CD4, and CD8, and the extracellular T cell activation markers, CD38 and HLA-DR as previously described (26). Briefly, PBMCs were incubated with fluorochrome-conjugated anti-CD3, -CD4, -CD8, -CD38 and -HLA-DR antibodies for 30 minutes at room temperature, washed, fixed with 1% formaldehyde, and analyzed by flow cytometry (LSR-II, BD Biosciences) using Diva software. All antibodies were obtained from BioLegend (San Diego, CA, USA).

### 5.5. Peptides

All peptides were synthesized by standard 9-fluorenylmethyloxycarbonyl (FMOC) chemistry and purified by reversed-phase high-performance liquid chromatography (purity at least 80%, usually >95%), mass spectrometry validated and lyophilized (Schafer-N, Copenhagen, Denmark). An overlapping peptide library systematically covering the entire proteome of the vaccine strain, YF-17D-204 (UniProt# P03314), was synthesized. This encompassed the entire YF precursor protein of 3411 aa, which could be represented by 850 peptides, each 15 aa long and overlapping its neighboring peptides by 11 aa. In addition, 50 peptides, which were predicted to be binders to HLA-A or -B supertype representatives by our NetMHCpan predictor, were selected from putative alternative translation initiation codon products (65). One hundred and seven (107, or 11.9%) of the total of 900 selected peptides were difficult to synthesize and/or purify; many of which had long stretches of hydrophobic aa. Adding one or two lysine to their N-or C-termini allowed the successful synthesis and purification of 77 of these “difficult to synthesize” peptides leaving 30 peptides that could not be synthesized and/or purified. Thus, 870 (97%) of the selected peptides could be included in this epitope screening effort. These peptides were initially organized in a 30×30 matrix, and eventually in four 15×15 matrices (**Supplementary Figure S3**).

### 5.6. Ex-vivo ELISpot assay

Fresh or thawed PBMCs were tested using an Interferon-γ (INFγ) specific ELI Spot assay as previously described (66). Briefly, 2-3 × 10^5^ cells/well were plated in an ELI Spot plate (MAHAS4510, Merck Millipore, USA) and *in vitro* cultured for 18-24 hours in media supplemented with or without peptide at 0.5 µM (or, as positive control, with 1 µg/ml Staphylococcal enterotoxin B (SEB, Sigma Aldrich, St. Louis, USA)). An AP conjugate substrate kit (Bio-Rad) was used for visualization of spots. ELI spots were counted using a CTL ImmunoSpot series 5 UV Analyzer. ImmunoSpot 5.0.9 software (C.T.L., Shaker Heights, USA) was used for analysis. Wells with spot-forming units SPU > 2 times the background wells were considered positive.

### 5.7. Cell culture and in vitro peptide stimulation

PBMCs were incubated overnight (37°C, 5% CO_2_) in X-vivo 15 media (Fisher Scientific) supplemented with 5% AB Serum (Invitrogen) and a mixture of relevant peptides at a concentration of 0.5µM of each peptide. The cells were harvested and washed, and subsequently plated in 24-well plates at a concentration of 5 × 10^6^/ml supplemented with 50U/ml IL-2 for expansion. Fresh media and IL-2 were supplemented every second day until the cells were harvested at day 8, and IL-15 (15ng/ml) was added the last four days.

### 5.8. *In vitro* intracellular cytokine staining (ICS) assay

*In vitro* cultured PBMC’s were harvested, washed, and aliquoted at 2-4 × 10^5^ cells/well. The cells were incubated with relevant peptide matrix column and row mixes (1uM/peptide) or single peptide (0.8 uM) for 4 h at 37°C, 5% CO_2_. Brefeldin A was added for the last 3 h of incubation. The cells were subsequently permeabilized (Becton Dickinson Permeabilizing solution 2) and stained with anti-CD3, -CD4, -CD8, -CD69, and -IFNγ, according to the “FastImmune” protocol (Becton Dickinson). The cells were subsequently analyzed by flow cytometry using a LSRII (BD Biosciences). Staining of more than 0.8% of CD8^+^ or CD4^+^ T cells was considered positive.

### 5.9. Peptide-HLA-I tetramers

HLA class I tetramers were produced as previously described (67). Briefly, recombinant, biotinylated HLA class I heavy chain, human β_2_-microglobulin and peptide were incubated in 50mM tris-maleate pH 6.8 and 0.1% Pluronic F68 for 48h at 18°C. The resulting monomers were tetramerized by addition of fluorochrome labelled Streptavidin (Streptavidin-phycoerythrin (SA-PE), Streptavidin-allophycocyanin (SA-APC), Streptavidin-Brilliant Violet 421 (SA-BV421), and/or Streptavidin-Brilliant Violet 605 (SA-BV605); all from BioLegend) sequentially over 60 min at a 1:4 molar ratio of streptavidin to monomer. For T cell analysis, pellets of 10^6^ PBMCs obtained *ex vivo*, or pellets from 2×10^5^ cells obtained from *in vitro* peptide stimulated cell cultures, were re-suspended in a 10µl tetramer solution at a final concentration of ≈ 30nM, and incubated for 20 min at room temperature, followed by 30 min incubation with fluorochrome conjugated anti-CD3, -CD8, -CD38 and -HLA-DR antibodies. The cells were analyzed by flowcytometry (Fortessa or LSR-II, BD Biosciences) using Diva software. Supplementary figure S4, a NS5_286-295_-A*01:01 tetramer *ex vivo* staining pre- and post-YF vaccination, illustrates the tetramer staining and background level.

### 5.10. Peptide-HLA-II tetramers

HLA-DRA and HLA-DRB chains were produced as previously described (43). For tetramer production, HLA-DRA and HLA-DRB chains were mixed in a 1:1.5 ratio and incubated in 3 µM peptides in PBS (pH 7.4) with 20% glycerol and 0.1% Pluronic F68 for 96h at 18°C. The resulting monomers were buffer changed into PBS with 5% glycerol and concentrated on 10kD Vivaspin (Satorius) and quantitated by Luminescent Oxygen Channeling Immunoassay (LOCI)-driven assay (43). The resulting monomers were tetramerized by addition of fluorochrome labelled Streptavidin (Streptavidin-phycoerythrin (SA-PE) or Streptavidin-allophycocyanin (SA-APC); both from BioLegend) sequentially over 60 min at a 1:4 molar ratio of streptavidin to monomer. For T cell analysis, pellets of 4×10^5^ cells obtained from *in vitro* peptide stimulated cell cultures, were re-suspended in a 40 µl tetramer diluted in media to a final concentration of ≈ 30 nM, and incubated for 1h at 37°C, followed by 30 min incubation with fluorochrome conjugated anti-CD3, -CD8, -CD38 and -HLA-DR antibodies. The cells were analyzed by flowcytometry (Fortessa or LSR-II, BD Biosciences) using Diva software.

### 5.11. Predictions of CD8^+^ T cell epitopes and HLA-I restriction

For each donor, all 15mer peptides eliciting a CD8^+^ T cell response were submitted to our bioinformatics predictor, NetMHCpan (37), which returned a prioritized list of predicted optimal epitopes, which could bind to any of the up to six HLA-A, -B, or-C molecules of the donor in question.

### 5.12. Predictions of CD4^+^ T cell epitopes and HLA-II restriction

For each donor, all 15mer peptides eliciting a CD4^+^ T cell response were submitted to our bioinformatics predictor, NetMHCIIpan (68), which returned a prioritized list of predicted epitopes including a predicted core-region, which could bind to any of the up to four HLA-DRB1, or-DRB3/4/5 molecules of the donor in question.

### 5.13. Peptide-HLA class I stability measurements

The stability of peptide-HLA class I complexes was measured using dissociation of ^125^I radiolabelled β_2_m in a scintillation proximity assay (SPA) as previously described (69). Briefly, recombinant, biotinylated HLA class I heavy chains were diluted into a refolding buffer containing the test peptide and trace amounts of ^125^I radiolabeled β_2_m, and allowed to refold at 18°C for 24 h in a Streptavidin-coated scintillation microplate (Flashplate PLUS, Perkin Elmer, Boston, MA). Dissociation was initiated by adding excess unlabeled β_2_m and placing the microplate in a scintillation counter (TopCount NXT, Packard) adjusted to 37°C. Reading the microplate continuously for 24 h allowed determination of the dissociation of radiolabeled β_2_m.

### 5.14. Biochemical peptide HLA class II binding assays

Peptide-HLA-II binding affinities were determined using a previously described Luminescent Oxygen Channeling Immunoassay (LOCI)-driven assay (43). Briefly, denatured and purified recombinant HLA-II alpha and beta chains were diluted into a refolding buffer (tris-maleate buffer, pH 6.6) with graded concentrations of the test peptide, and incubated for 48 h at 18°C to allow for equilibrium to be reached. The peptide concentration leading to half-saturation (ED_50_) was determined as previously described (43). Under the limited receptor concentrations used here, the ED_50_ reflects the affinity of the interaction.

### 5.15. Statistics

GraphPad Prism 8 was used for statistical analyses (unpaired and paired Mann-Whitney-Wilcoxon tests, unpaired and paired t-tests, Fishers exact test, and ROC analysis).

## 7. Conflict of Interest

*The authors declare that the research was conducted in the absence of any commercial or financial relationships that could be construed as a potential conflict of interest*.

## 8. Author Contributions

AS, JPC, MN, ART and SB contributed to the conception and design of the study; SB, AS, ST, MG, and MBH recruited donors, vaccinated and collected blood samples before and after vaccination; MK, MR, MNH performed biochemical analysis; AS, MN and SB performed the statistical analysis; AS wrote the first draft of the manuscript; AS and SB wrote the manuscript with sections contributed by MN and ART. All authors contributed to manuscript revision, read and approved the submitted version.

## 9. Funding

This project has been funded in part with Federal funds from the National Institute of Allergy and Infectious Diseases, National Institutes of Health, Department of Health and Human Services, under, Contract No. HHSN272200900045C

## 10. Acknowledgments

The authors thank doctors and nurses at the two recruiting centers and the Blood Bank of the University Hospital, and members of the Buus laboratory for expert technical assistance.

