## Supplemental material for "A systematic, unbiased mapping of CD8^+^ and CD4^+^ T cell epitopes in Yellow Fever vaccinees"

**Supplementary results and discussion of the performance of the hybrid forward-reverse immunology (HFRI) vs reverse immunology approaches.**

Additional opportunities arose for comparing the HFRI and reverse immunology approaches. For donor YF1067, the 27 stimulatory 15mer peptides were submitted to the contemporary NetMHCpan 2.4, which returned 156 submer-HLA-I predictions per stimulatory peptide. For each stimulatory peptide, we retrospectively asked how each of the 19 validated epitopes of donor YF1067 ranked amongst the 156 submers predictions. In 23(≈85%) of the 27 submissions, the number one ranking prediction ended up being validated as an epitope (**Table I**). If one applied the contemporary NetMHCpan 2.4 predictor at a stringent %Rank cut-off of 0.5%, it would have correctly identified 17 (89%) of the 19 epitopes (true positives); missed 2 (false negatives), and erroneously included 14 (false positives) leading to a false positive rate of 14/(14+19) = 42%. The current version, NetMHCpan4.0, could have been applied at an even more stringent %Rank cut-off of 0.25%, which would have correctly identified 18 (95%) of the 19 epitopes (true positives); 1 would have been missed (false negative), and 6 would have been erroneously included (false positives) leading to a false positive rate of 6/(6+19) = 24%.

Had the predictors been applied, not to the 27 stimulatory peptides, but to the entire YFV proteome, with the above cut-offs and the HLA-A and -B allotypes of YF1067, the number of false positive predictions included by NetMHCpan 2.4 would have been 187, whereas the current NetMHCpan4.0 would have included 158 false positive predictions.

This supports the general conclusion that the present HFRI approach ranks epitope at the very top of the list of candidates while decimating the false discovery rate. It also demonstrates that the current predictors have improved compared to older versions maintaining a high sensitivity while reducing the false positive rate. Nonetheless, there is still room for further improvements.

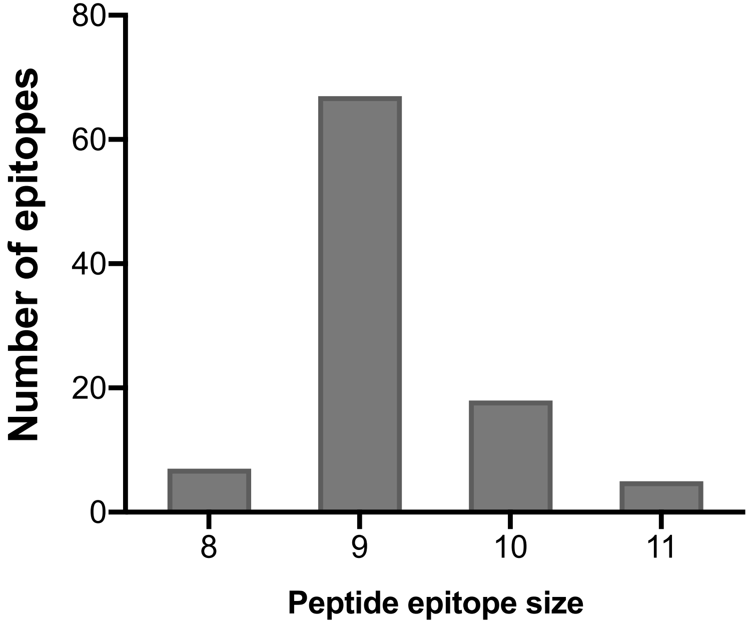


Supplementary Figure S1. The length distribution of the identified CD8+ T cell epitopes.
Shown is the number of identified YFV-specific, HLA-class I restricted CD8+ T cell epitopes of 8-, 9-, 10- and 11-mer size for all tetramer validated epitopes.

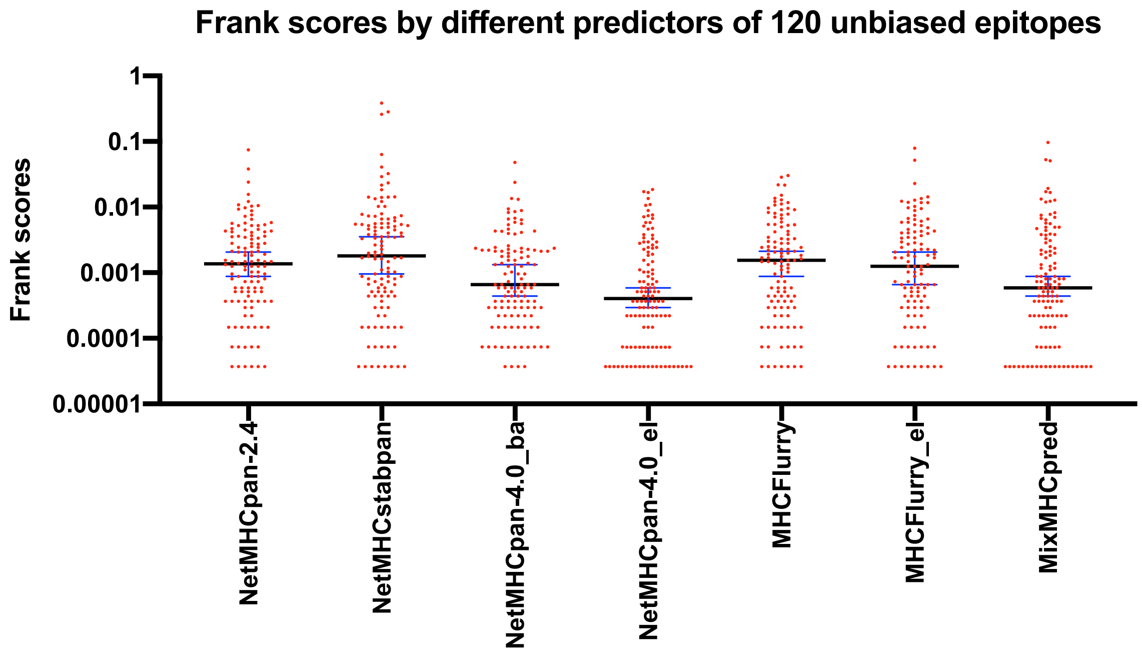


| | | | | | | | |
|---|---|---|---|---|---|---|---|
| AUC | 0.99636 | 0.98743 | 0.99755 | 0.99797 | 0.99631 | 0.99604 | 0.99562 |
| Median Frank score | 0.00136 | 0.00180 | 0.00066 | 0.00040 | 0.00154 | 0.00125 | 0.00059 |
| Wilcoxon matched-pairs signed rank test. p-value | 0.0001 | <0.0001 | 0.0102 | NA | <0.0001 | <0.0001 | 0.0156 |

Supplementary Figure S2. The Yellow Fever proteome of 3411 aa is submitted to the various indicated predictors along with the HLA-I restriction elements used by the epitopes discovered by the HFRI approach, and the bindings of all 13610 possible 8-11mer peptides per HLA-I are predicted. Per predictor, the median Frank scores (the frequencies of False Positive predictions) of the 120 HFRI discoverted epitopes is determined. The median Frank is indicated by black bars and the +/- 95% CI is indicated by blue error bars. For illstrative purposes, the perfect predictions with a Frank score of 0 has been recoded to a Frank score of 0.000037 and are clearly visible as the bottom line of symbols.

Supplementary Figure S3. Using peptide matrices to screeen for peptides containing YFV-derived T cell epitopes

A

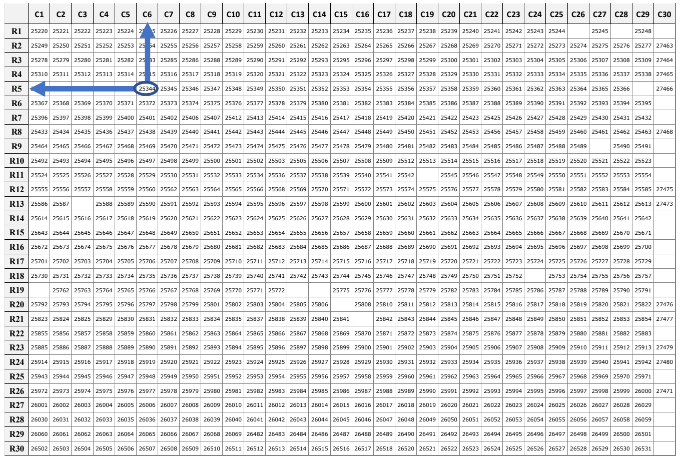

B

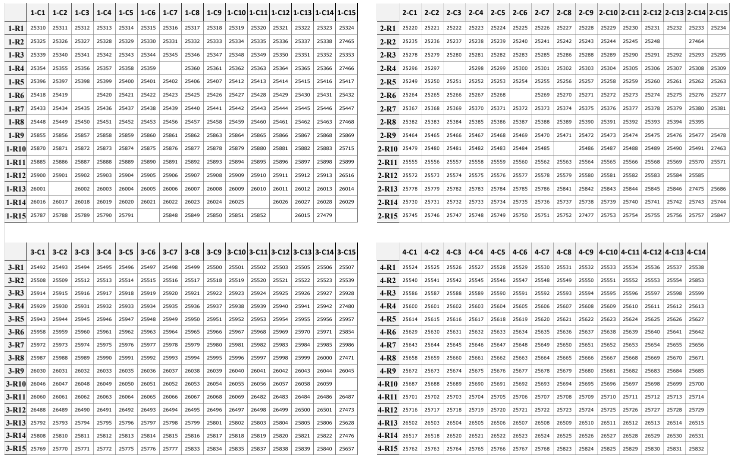

A) The 30x30 matrix approach. The 870 Yellow Fever peptides were organized into a 30x30 matrix. Row pools, R1-R30, contained up to 30 different peptides pooled row-wise; column pools, C1-C30, contained up to 30 different peptides pooled column-wise. Every single peptide was therefore present in one specific column and one specific row pool (the oval exemplifies one peptide, 25344, which was present in both column 6 and row 5). Thus, every intersection of T cell stimulatory column and row pools tentatively identified a T cell epitope-containing peptide for further deconvolution and study.

B) The 4x(15x15) matrices approach. The 870 Yellow Fever peptides were organized into four 15x15 matrices. Following the same general pooling strategy as for the 30x30 matrix, the 4x(15x15) matrix strategy led to the generation of 4x15 column pools and 4x15 row pools (actually, matrix 4 only contained 14 columns). Per submatrix, every intersection of T cell stimulatory column and row pools tentatively identified a T cell epitope-containing peptide for further deconvolution and study

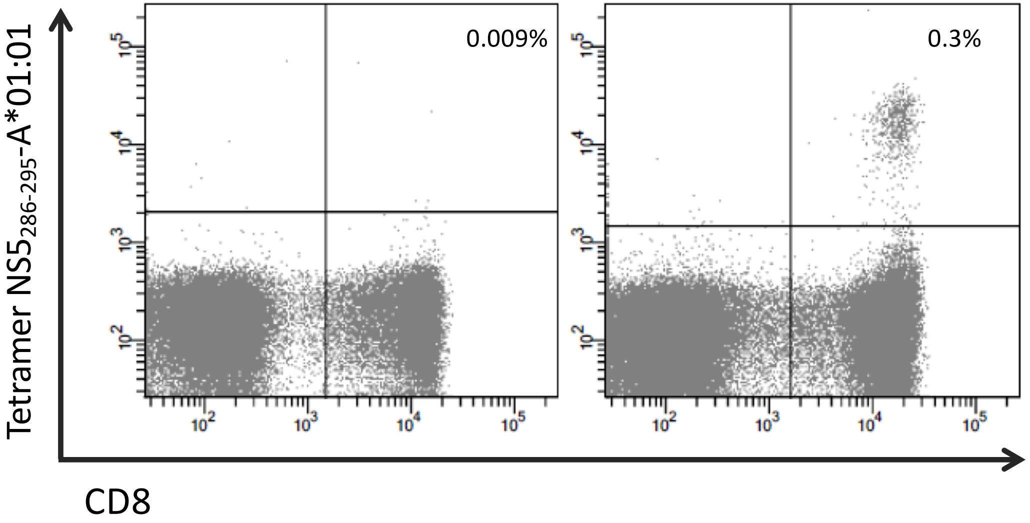


Supplementary Figure S4: HLA-A*01:01 positive donor stained for the dominant HLA-A1*01:01 epitope, NS5(286-295). PBMCs from the donor before vaccination (left plot) and after Yellow Fever vaccination (right plot) were stained with anti CD3-Pacific Blue, CD8-PerCp, and PE labeled NS5(286-295)/HLA-A1*01:01 tetramer. Shown is CD3+ gated T cells and the percentage given is Tetramer+CD8+ T cells.

| Proteom position | Protein position | Sequence | HLA restriction | Predicted affinity (%RANK) | Stability ($t_{½}$; h) | Responders /donor tested | Tetramer validated (%CD8$^{+}$ T cells) | Reference |
|---|---|---|---|---|---|---|---|---|
| YFV_pp61-69 | CapsidC (61-69) | ITAHLKRLW | B*57:01 | 0.05 | 9.0 | 3/11 | 0.01-0.02 | |
| YFV_pp64-72 | CapsidC (64-72) | HLKRLWKML | B*08:01 | 0.30 | 0.5 | 8/42 | 0.03-0.30 | |
| YFV_pp102-110 | ER anchor (1-9) | SHDVLTVQF | B*38:01 | 0.40 | 1.6 | 7/7 | 0.02-0.20 | |
| YFV_pp103-110 | ER anchor (2-9) | HDVLTVQF | B*37:01 | 3.00 | 19.4 | 2/2 | 0.02-0.50 | |
| YFV_pp331-339 | EnvE (46-54) | ETVAIDRPA | A*68:02 | 0.18 | 15.8 | 2/2 | 0.10-0.40 | |
| YFV_pp453-460 | EnvE (168-175) | QEVEFIGY | B*18:01 | 0.08 | 4.1 | 13/14 | 0.01-0.70 | |
| YFV_pp471-479 | EnvE (186-194) | TAVDFGNSY | B*35:01 | 0.25 | 2.7 | 9/23 | 0.01-0.10 | |
| YFV_pp485-494 | EnvE (200-209) | TESWIVDRQW | B*44:02 | 0.15 | NA | 33/35 | 0.01-1.50 | (PMID:28386132)[1] |
| YFV_pp485-494 | EnvE (200-209) | TESWIVDRQW | B*44:03 | 0.13 | NA | 8/12 | 0.03-0.20 | |
| YFV_pp486-494 | EnvE (201-209) | ESWIVDRQW | B*57:01 | 0.30 | 1.9 | 13/15 | 0.01-0.20 | |
| YFV_pp492-500 | EnvE (207-215) | RQWAQDLTL | B*40:02 | 0.40 | 2.4 | 2/7 | 0.01-0.20 | |
| YFV_pp492-500 | EnvE (207-215) | RQWAQDLTL | B*13:02 | 0.40 | 2.4 | 4/4 | 0.05-0.30 | (PMID:28386132)[1] |
| YFV_pp500-510 | EnvE (215-225) | LPWQSGSGGVW | B*53:01 | 0.03 | NA | 3/3 | 0.20-0.30 | |
| YFV_pp500-510 | EnvE (215-225) | LPWQSGSGGVW | B*35:01 | 0.20 | 2.9 | 12/22 | 0.01-0.20 | |
| YFV_pp511-519 | EnvE (226-234) | REMHHLVEF | B*37:01 | 0.03 | 19.8 | 3/3 | 0.01-0.30 | |
| YFV_pp511-519 | EnvE (226-234) | REMHHLVEF | B*40:01 | 0.05 | 1.5 | 19/30 | 0.01-0.80 | |
| YFV_pp517-525 | EnvE (232-240) | VEFEPPHAA | B*40:02 | 0.80 | 12.1 | 6/7 | 0.01-0.10 | |
| YFV_pp517-525 | EnvE (232-240) | VEFEPPHAA | B*50:01 | 0.10 | 3.8 | 2/3 | 0.10-0.20 | |
| YFV_pp569-577 | EnvE (284-292) | RVKLSALTL | B*07:02 | 1.50 | 4.8 | 5/41 | 0.03-0.40 | (PMID: 22039500)[2] |
| YFV_pp569-578 | EnvE (284-293) | RVKLSALTLK | A*03:01 | 0.25 | 36.3 | 29/39 | 0.01-0.20 | |
| YFV_pp651-660 | EnvE (366-375) | EVNPPFGDSY | A*25:01 | 0.13 | 3.1 | 2/11 | 0.03-0.10 | |
| YFV_pp651-660 | EnvE (366-375) | EVNPPFGDSY | A*26:01 | 0.13 | 1.5 | 3/10 | 0.02-0.03 | |
| YFV_pp729-737 | EnvE (444-452) | GLFGGLNWI | A*02:01 | 0.30 | 5.1 | 12/89 | 0.01-0.10 | (PMID: 22039500)[2] |
| YFV_pp759-767 | EnvE (474-482) | SMSMILVGV | A*02:01 | 0.30 | 16.4 | 27/95 | 0.01-0.10 | |
| YFV_pp802-810 | NS1 (24-32) | DSDDWLNKY | A*01:01 | 0.05 | 1.4 | 40/53 | 0.01-0.20 | |
| YFV_pp883-891 | NS1 (105-113) | SRIRDGLQY | B*27:02 | 0.80 | 9.0 | 1/1 | 0.05-0.05 | |
| YFV_pp884-893 | NS1 (106-115) | RIRDGLQYGW | A*32:01 | 0.40 | 107.9 | 15/18 | 0.02-0.10 | |
| YFV_pp884-893 | NS1 (106-115) | RIRDGLQYGW | B*57:01 | 0.40 | 6.3 | 9/15 | 0.01-0.10 | |
| YFV_pp885-893 | NS1 (107-115) | IRDGLQYGW | B*27:02 | 1.00 | 2.4 | 1/1 | 0.40-0.40 | |
| YFV_pp894-902 | NS1 (116-124) | KTWGKNLVF | A*32:01 | 0.03 | 32.1 | 19/19 | 0.10-2.80 | (PMID:28386132)[1] |
| YFV_pp894-902 | NS1 (116-124) | KTWGKNLVF | B*58:01 | 0.25 | 9.3 | 3/3 | 0.20-0.40 | |
| YFV_pp894-902 | NS1 (116-124) | KTWGKNLVF | B*57:01 | 0.10 | 5.3 | 14/15 | 0.02-0.80 | (PMID:25674793)[1] |
| YFV_pp945-953 | NS1 (167-175) | VYMDAVFEY | A*29:02 | 0.08 | 39.1 | 3/9 | 0.03-0.10 | |
| YFV_pp945-953 | NS1 (167-175) | VYMDAVFEY | A*24:02 | 0.80 | 32.8 | 7/33 | 0.01-0.20 | (PMID 23338234)[3] |
| YFV_pp970-978 | NS1 (192-200) | KSAHGSPTF | B*58:01 | 0.13 | 13.7 | 2/3 | 0.10-0.10 | |
| YFV_pp983-991 | NS1 (205-213) | HEVNGTWMI | B*40:01 | 0.13 | 1.2 | 2/31 | 0.02-0.03 | |
| YFV_pp1002-1010 | NS1 (224-232) | CEWPLTHTI | B*49:01 | 0.01 | 11.7 | 2/3 | 0.05-0.05 | |
| YFV_pp1002-1010 | NS1 (224-232) | CEWPLTHTI | B*40:01 | 0.40 | 0.9 | 6/26 | 0.02-0.80 | |
| YFV_pp1114-1123 | NS1 (336-345) | RPRKTHESHL | B*07:02 | 0.03 | 4.8 | 19/44 | 0.01-0.20 | |
| YFV_pp1114-1124 | NS1 (336-346) | RPRKTHESHLV | B*07:02 | 0.10 | 3.0 | 12/44 | 0.01-0.20 | |
| YFV_pp1119-1127 | NS1 (341-349) | HESHLVRSW | B*44:02 | 0.05 | NA | 30/33 | 0.02-0.40 | |
| YFV_pp1119-1127 | NS1 (341-349) | HESHLVRSW | B*44:03 | 0.05 | NA | 9/13 | 0.10-0.30 | |
| YFV_pp1131-1140 | NS2A (1-10) | GEIHAVPFGL | B*40:01 | 0.01 | 1.5 | 22/28 | 0.01-0.50 | |
| YFV_pp1131-1141 | NS2A (1-11) | GEIHAVPFGLV | B*40:01 | 0.25 | 1.2 | 13/31 | 0.01-0.30 | |
| YFV_pp1134-1143 | NS2A (4-13) | HAVPFGLVSM | B*35:08 | 0.80 | 13.5 | 2/2 | 1.30-9.70 | (PMID: 11853408)[4]; (PMID:19740333)[5] |
| YFV_pp1134-1143 | NS2A (4-13) | HAVPFGLVSM | B*35:03 | 0.80 | 7.7 | 6/6 | 0.20-4.50 | (PMID: 11853408)[4]; (PMID:19740333)[5] |
| YFV_pp1134-1143 | NS2A (4-13) | HAVPFGLVSM | B*35:01 | 0.40 | 7.0 | 22/24 | 0.10-1.70 | (PMID: 11853408)[4]; (PMID:19740333)[5]; (PMID:28386132)[1] |
| YFV_pp1200-1207 | NS2A (70-77) | DAMYMALI | B*51:01 | 0.80 | 17.3 | 13/18 | 0.03-2.20 | |
| YFV_pp1227-1234 | NS2A (97-104) | SPRERLVL | B*07:02 | 0.01 | 4.1 | 4/44 | 0.01-0.02 | |
| YFV_pp1227-1236 | NS2A (97-106) | SPRERLVLTL | B*07:02 | 0.03 | 1.8 | 40/44 | 0.02-4.00 | |
| YFV_pp1317-1325 | NS2A (187-195) | SMQKTIPLV | A*02:01 | 0.80 | 27.3 | 43/95 | 0.01-0.10 | |
| YFV_pp1397-1405 | NS2B (43-51) | SVAGRVDGL | A*02:05 | 4.00 | 18.7 | 5/5 | 0.02-0.40 | |
| YFV_pp1464-1472 | NS2B (110-118) | HPFALLLVL | B*35:01 | 0.30 | 5.9 | 22/23 | 0.05-0.90 | (PMID: 11853408)[4]; (PMID:19740333)[5]; (PMID:28386132)[1] |
| YFV_pp1464-1472 | NS2B (110-118) | HPFALLLVL | B*35:03 | 0.13 | 5.9 | 5/6 | 0.20-0.80 | (PMID: 11853408)[4]; (PMID:19740333)[5]; (PMID:28386132)[1] |
| YFV_pp1471-1479 | NS2B (117-125) | VLAGWLFHV | A*02:01 | 0.01 | 27.7 | 75/86 | 0.01-0.50 | (PMID: 22039500)[2] |
| YFV_pp1488-1496 | NS3 (4-12) | VLWDIPTPK | A*11:01 | 0.50 | 9.0 | 16/28 | 0.01-0.20 | |
| YFV_pp1488-1496 | NS3 (4-12) | VLWDIPTPK | A*03:01 | 0.18 | 6.2 | 31/40 | 0.01-0.40 | (PMID:28386132)[1] |
| YFV_pp1508-1516 | NS3 (24-32) | IYGIFQSTF | A*24:02 | 0.08 | 40.7 | 22/33 | 0.01-1.00 | (PMID:28386132)[1] |
| YFV_pp1508-1516 | NS3 (24-32) | IYGIFQSTF | A*23:01 | 0.13 | 40.6 | 4/5 | 0.01-0.20 | |
| YFV_pp1557-1565 | NS3 (73-81) | SVKEDLVAY | B*15:01 | 0.80 | 14.5 | 28/30 | 0.03-1.10 | (PMID:28386132)[1] |
| YFV_pp1557-1565 | NS3 (73-81) | SVKEDLVAY | B*35:01 | 1.50 | 3.2 | 13/23 | 0.01-0.10 | (PMID:28386132)[1] |
| YFV_pp1608-1615 | NS3 (124-131) | GEIGAVAL | B*40:01 | 0.01 | 3.2 | 5/17 | 0.02-0.10 | |

| | | | | | | | | |
|---|---|---|---|---|---|---|---|---|
| YFV_pp1632-1641 | NS3 (148-157) | EVIGLYGNGI | A*68:02 | 0.50 | 2.9 | 1/2 | 0.04-0.04 | |
| YFV_pp1635-1643 | NS3 (151-159) | GLYGNGILV | A*02:01 | 1.50 | 20.1 | 27/80 | 0.01-0.20 | (PMID: 22039500)[2] |
| YFV_pp1690-1697 | NS3 (206-213) | RRFLPQIL | C*06:02 | 0.15 | 1.4 | 6/28 | 0.10-2.78 | |
| YFV_pp1689-1697 | NS3 (205-213) | TRRFLPQIL | C*06:02 | 8.00 | 17.7 | 27/28 | 0.02-1.90 | |
| YFV_pp1702-1710 | NS3 (218-226) | RRRLRTLVL | B*14:02 | 0.15 | 0.9 | 4/4 | 0.05-0.40 | (PMID: 18762226)[6] |
| YFV_pp1718-1727 | NS3 (234-243) | SEMKEAFHGL | B*40:01 | 0.05 | 1.3 | 16/29 | 0.01-0.40 | |
| YFV_pp1728-1736 | NS3 (244-252) | DVKFHTQAF | A*25:01 | 0.15 | 3.2 | 10/11 | 0.02-0.30 | (PMID: 18762226)[6] |
| YFV_pp1770-1778 | NS3 (286-294) | IIMDEAHFL | A*02:05 | 0.20 | 20.7 | 5/5 | 0.10-0.70 | |
| YFV_pp1770-1778 | NS3 (286-294) | IIMDEAHFL | A*02:01 | 0.50 | 9.2 | 43/93 | 0.01-0.10 | (PMID: 22039500)[2] |
| YFV_pp1777-1785 | NS3 (293-301) | FLDPASIAA | A*02:01 | 0.40 | 9.9 | 56/84 | 0.01-0.70 | |
| YFV_pp1831-1839 | NS3 (347-355) | EPWNTGHDW | B*53:01 | 0.10 | NA | 2/3 | 0.10-0.10 | |
| YFV_pp1910-1918 | NS3 (426-434) | RVLDCRTAF | A*32:01 | 0.80 | 31.1 | 4/19 | 0.05-0.10 | |
| YFV_pp1991-1999 | NS3 (507-515) | GMVAPLYGV | A*02:01 | 0.10 | 9.4 | 37/96 | 0.01-0.20 | |
| YFV_pp2030-2039 | NS3 (546-555) | LPVWLSWQVA | B*56:01 | 0.05 | 2.3 | 3/3 | 0.10-0.10 | |
| YFV_pp2060-2068 | NS3 (576-584) | ILNDSGETV | A*02:01 | 1.50 | 12.2 | 42/92 | 0.01-0.40 | |
| YFV_pp2128-2136 | NS4A (21-29) | GEAMDTISV | B*40:01 | 0.40 | 1.3 | 7/32 | 0.01-0.20 | |
| YFV_pp2129-2137 | NS4A (22-30) | EAMDTISVF | B*35:01 | 0.10 | 3.7 | 3/21 | 0.01-0.03 | |
| YFV_pp2130-2138 | NS4A (23-31) | AMDTISVFL | A*02:01 | 0.80 | 11.0 | 5/95 | 0.01-0.03 | (PMID:27017899) |
| YFV_pp2152-2160 | NS4A (45-53) | SMMPEAMTI | B*52:01 | 0.10 | 9.1 | 3/3 | 0.01-0.03 | |
| YFV_pp2154-2163 | NS4A (47-56) | MPEAMTIVML | B*35:03 | 0.05 | 3.9 | 5/6 | 0.10-0.80 | |
| YFV_pp2154-2163 | NS4A (47-56) | MPEAMTIVML | B*35:01 | 0.30 | 3.8 | 21/25 | 0.03-1.00 | |
| YFV_pp2154-2163 | NS4A (47-56) | MPEAMTIVML | B*53:01 | 0.15 | 2.9 | 2/3 | 0.10-0.10 | |
| YFV_pp2155-2163 | NS4A (48-56) | PEAMTIVML | B*40:01 | 1.00 | ND | 13/17 | 0.10-0.80 | |
| YFV_pp2180-2189 | NS4A (73-82) | SPKGISRMSM | B*07:02 | 0.10 | 8.6 | 29/43 | 0.01-0.10 | |
| YFV_pp2389-2397 | NS4B (133-141) | KLAQRRVFH | A*03:01 | 1.00 | 8.6 | 26/32 | 0.01-1.40 | |
| YFV_pp2421-2429 | NS4B (165-173) | ALYEKKLAL | A*02:01 | 2.00 | 19.2 | 3/85 | 0.01-0.50 | |
| YFV_pp2421-2429 | NS4B (165-173) | ALYEKKLAL | B*08:01 | 0.25 | 0.4 | 28/41 | 0.01-0.20 | |
| YFV_pp2423-2431 | NS4B (167-175) | YEKKLALYL | B*40:02 | 0.40 | 24.4 | 7/7 | 0.04-0.60 | |
| YFV_pp2423-2431 | NS4B (167-175) | YEKKLALYL | B*40:01 | 0.30 | 1.1 | 4/28 | 0.03-0.30 | |
| YFV_pp2470-2478 | NS4B (214-222) | LLWNGPMAV | A*02:01 | 0.13 | 38.2 | 98/98 | 0.03-11.50 | (PMID 19933869) |
| YFV_pp2494-2502 | NS4B (238-246) | VMYNLWKMK | A*03:01 | 0.05 | 0.8 | 17/38 | 0.01-2.40 | |
| YFV_pp2524-2533 | NS5 (18-27) | LLDKRQFELY | A*01:01 | 0.05 | 11.7 | 32/52 | 0.01-0.20 | |
| YFV_pp2571-2579 | NS5 (65-73) | FHERGYVKL | B*38:01 | 0.10 | 5.2 | 7/7 | 0.10-0.60 | |
| YFV_pp2571-2579 | NS5 (65-73) | FHERGYVKL | B*39:01 | 0.15 | 0.6 | 5/6 | 0.10-1.00 | |
| YFV_pp2669-2677 | NS5 (163-171) | RVLDTVEKW | A*32:01 | 0.80 | 101.6 | 11/19 | 0.01-0.10 | |
| YFV_pp2669-2677 | NS5 (163-171) | RVLDTVEKW | B*58:01 | 0.13 | 32.9 | 2/3 | 0.10-0.40 | |
| YFV_pp2669-2677 | NS5 (163-171) | RVLDTVEKW | B*57:01 | 0.15 | 16.2 | 13/15 | 0.01-0.20 | |
| YFV_pp2706-2714 | NS5 (200-208) | RRFGGTVIR | B*27:05 | 0.05 | 22.9 | 6/12 | 0.02-0.10 | |
| YFV_pp2711-2719 | NS5 (205-213) | TVIRNPLSR | A*68:01 | 0.50 | 11.9 | 11/19 | 0.03-0.70 | |
| YFV_pp2723-2731 | NS5 (217-225) | HEMYYVSGA | B*50:01 | 0.08 | NA | 3/3 | 0.04-0.10 | |
| YFV_pp2762-2770 | NS5 (256-264) | DVILPIGTR | A*68:01 | 0.50 | 37.4 | 20/20 | 0.01-0.90 | |
| YFV_pp2792-2801 | NS5 (286-295) | KSEYMTSWFY | A*01:01 | 0.08 | 44.2 | 50/52 | 0.01-4.60 | (PMID:28386132)[1] |
| YFV_pp2854-2862 | NS5 (348-356) | TPFGQQRVF | B*35:01 | 0.40 | 3.6 | 8/22 | 0.01-0.10 | |
| YFV_pp2879-2887 | NS5 (373-381) | KIMKVVNRW | B*58:01 | 0.18 | 19.2 | 1/3 | 0.01-0.01 | |
| YFV_pp2882-2890 | NS5 (376-384) | KVVNRWLFR | A*03:01 | 0.40 | 6.5 | 2/38 | 0.02-0.03 | |
| YFV_pp2974-2982 | NS5 (468-476) | KAKGSRAIW | B*57:01 | 0.13 | 11.2 | 10/15 | 0.01-0.10 | |
| YFV_pp2977-2985 | NS5 (471-479) | GSRAIWYMW | B*57:01 | 0.08 | 9.1 | 5/15 | 0.01-0.02 | |
| YFV_pp2981-2990 | NS5 (475-484) | IWYMWLGARY | A*29:02 | 0.10 | 0.6 | 2/8 | 0.10-0.20 | |
| YFV_pp2982-2990 | NS5 (476-484) | WYMWLGARY | A*29:02 | 0.25 | 24.7 | 8/9 | 0.10-0.50 | (PMID:28386132)[1] |
| YFV_pp2983-2990 | NS5 (477-484) | YMWLGARY | A*29:02 | 0.03 | 3.0 | 3/9 | 0.03-0.30 | |
| YFV_pp2983-2990 | NS5 (477-484) | YMWLGARY | A*01:01 | 1.50 | 67.3 | 47/52 | 0.01-0.80 | |
| YFV_pp3024-3032 | NS5 (518-526) | YVIRDLAAM | B*35:01 | 0.13 | 5.9 | 16/20 | 0.01-0.10 | |
| YFV_pp3063-3071 | NS5 (557-565) | YMSPHHKKL | A*02:05 | 1.50 | 16.9 | 4/5 | 0.10-0.20 | |
| YFV_pp3178-3186 | NS5 (672-680) | RPIDDRFGL | B*07:02 | 0.15 | 6.3 | 45/45 | 0.01-0.80 | (PMID: 23338234)[3]; (PMID: 28386132)[1] |
| YFV_pp3178-3187 | NS5 (672-681) | RPIDDRFGLA | B*07:02 | 0.80 | 3.1 | 8/46 | 0.01-0.50 | |
| YFV_pp3178-3188 | NS5 (672-682) | RPIDDRFGLAL | B*07:02 | 0.03 | 2.5 | 41/47 | 0.01-0.60 | |
| YFV_pp3178-3188 | NS5 (672-682) | RPIDDRFGLAL | B*35:03 | 0.25 | 3.5 | 3/6 | 0.01-0.10 | |
| YFV_pp3389-3399 | NS5 (883-893) | YTDYLTVMDRY | A*01:01 | 0.01 | 42.0 | 48/53 | 0.06-0.70 | |

**Table SI. A complete list of the 120 peptide-specific, HLA-I-restricted CD8+ T cell epitopes identified by the HFRI approach.**

The HFRI-based $CD8^+$ T cell epitope discovery process detailed for donor YF1067 was extended to 50 randomely selected, primary YFV-vaccinated donors. Ninety-two different epitopes restricted by 40 HLA-I allotypes were discovered. The peptide-HLA-I affinity was predicted by NetMHCpan 2.4 (given in %Rank; the lower, the better); the stability was measured (given as half-life at 37$^o$C in hours; the longer, the better). Productively interaction peptide-HLA-I combinations were used to design peptide-HLA-I tetramers. the resulting tetramers were used to stain and ex vivo analyze CD8+ T cells by flow cytometry gating on CD3+ CD8+ T cells. The prevalence of these responses is given as "the number of positive donors/the number of donors tested". The magnitudes are given as the range of frequencies (in %) of *ex vivo* tetramer-stained CD8+ T cells. Note, that for the prevalence and magnitude measurements, the specific responses identified in 50 vaccinees was extended to additional donors within our cohort of primary vaccinee, who expressed the relevant HLA-I allotype.
PubMed ID (PMID) references are given in the event that an epitope had been reported before this study.

| Proteome position | Protein position | Sequence | HLA restriction | Predicted affinity (%RANK) | Stability ($t_{½}$; h) | Responders per donors tested | Tetramer validated (%CD8$^{+}$ T cells) |
|---|---|---|---|---|---|---|---|
| YFV_pp438-446 | EnvE (153-161) | NTDIKTLKF | A*01:01 | 0.15 | 1.70 | 46/51 | 0.02-0.90 |
| YFV_pp511-519 | EnvE (226-234) | REMHHLVEF | B*40:02 | 0.03 | 17.7 | 3/7 | 0.01-0.02 |
| YFV_pp1131-1140 | NS2A (1-10) | GEIHAVPFGL | B*40:02 | 0.05 | 11.8 | 5/7 | 0.03-0.10 |
| YFV_pp1504-1512 | NS3 (20-28) | LEDGIYGIF | B*40:01 | 0.4 | 2.30 | 6/27 | 0.01-0.10 |
| YFV_pp2762-2770 | NS5 (256-264) | DVILPIGTR | A*33:01 | 0.18 | NA | 2/2 | 0.01-0.10 |
| YFV_pp3024-3032 | NS5 (518-526) | YVIRDLAAM | A*26:01 | 0.01 | 10.6 | 7/10 | 0.01-0.30 |
| YFV_pp3089-3098 | NS5 (583-592) | RPAPGGKAYM | B*07:02 | 0.03 | 8.5 | 22/44 | 0.01-0.20 |
| YFV_pp3116-3124 | NS5 (610-618) | ALNTITNLK | A*03:01 | 0.10 | 14.8 | 12/33 | 0.01-0.10 |
| YFV_pp3268-3276 | NS5 (762-770) | YANMWSLMY | B*35:01 | 0.30 | 8.6 | 7/22 | 0.01-0.10 |

**Supplementary Table SII. CD8+ T cell epitopes derived from past epitope discovery efforts.**

A panel of 533 YFV-derived peptides obtained from past epitope discovery efforts were evaluated by NetMHCpan 2.4 in an effort to discover additional CD8+ T cell epitopes. The stability of predicted peptide-HLA-I binders were verified experimentally. An additional 90 peptide-HLA-I tetramers could be generated and tested in appropriate primary YFV-vaccinated donors. Additionally 9 epitopes and their restriction elements were identified. The prevalence and response magnitude of the corresponding CD8+ T cell responses were evaluated as descibed in Supplementary Table SI.

| | | Predicted | Measured | T cell analysis | |
|---|---|---|---|---|---|
| Protein Position | Sequence | %Rank | Stability (T½, h) | Responders /donor tested | Suggested by HFRI |
| NS2A (97-104) | SPRERLVL | 0.01 | 4.1 | 4/44 | Yes[1] |
| NS1 (243-251) | MPRSIGGPV | 0.03 | 4.6 | 0 | |
| NS2A (97-106) | SPRERLVLTL | 0.03 | 1.8 | 40/44 | Yes[1] |
| NS5 (672-682) | RPIDDRFGLAL | 0.03 | 2.5 | 41/47 | Yes |
| NS1 (336-345) | RPRKTHESHL | 0.03 | 4.8 | 19/44 | Yes |
| NS5 (583-592) | RPAPGGKAYM | 0.03 | 8.5 | 22/44 | No |
| NS4B (163-173) | MPALYEKKLAL | 0.05 | 2.0 | 0/16 | |
| CapsidC (15-23) | MVRRGVRSL | 0.05 | 3.3 | 0/16 | |
| NS5 (112-119) | KPMNVQSL | 0.05 | 3.4 | 0/18 | |
| NS5 (246-253) | RPTGKVTL | 0.08 | 6.5 | 0/18 | |
| NS1 (336-346) | RPRKTHESHLV | 0.10 | 3.0 | 12/44 | Yes |
| NS4A (73-82) | SPKGISRMSM | 0.10 | 8.6 | 29/43 | Yes[2] |
| NS4A (47-55) | MPEAMTIVM | 0.10 | 0.8 | 0/17 | Yes[3] |
| NS4B (218-228) | GPMAVSMTGVM | 0.10 | 14.4 | 0/16 | |
| NS3 (133-142) | YPSGTSGSPI | 0.10 | 2.5 | 0/17 | |
| prM (128-136) | NPFFAVTAL | 0.12 | 2.3 | 0/18 | |
| NS3 (367-376) | LPSIRAANVM | 0.12 | 5.7 | 0/18 | |
| NS4B (107-117) | MPLLCGIGCAM | 0.12 | 3.5 | 0/18 | |
| NS1 (102-111) | HPFSRIRDGL | 0.12 | 3.0 | 0/16 | |
| NS5 (672-680) | RPIDDRFGL | 0.15 | 6.3 | 45/45 | Yes |
| NS5 (559-569) | SPHHKKLAQAV | 0.15 | 1.4 | 0/18 | |
| NS5 (259-266) | LPIGTRSV | 0.17 | 0.7 | 0/18 | Yes |
| NS3 (199-209) | HPGAGKTRRFL | 0.20 | 1.6 | 0/17 | |
| NS2B (110-118) | HPFALLLVL | 0.20 | NF | ND | |
| NS5 (710-720) | VPFCSHHFHEL | 0.25 | 3.9 | 0/18 | |
| NS4A (100-108) | KPTHISYVM | 0.25 | 6.5 | 0/18 | |
| NS1 (266-273) | GPWMQVPL | 0.25 | 0.4 | 0/18 | |
| NS5 (185-192) | APYMPDVL | 0.25 | 6.4 | 0/18 | |
| NS3 (361-370) | RPTAWFLPSI | 0.25 | 1.7 | 0/17 | |
| NS1 (226-236) | WPLTHTIGTSV | 0.25 | 2.6 | 0/18 | |
| NS5 (583-590) | RPAPGGKA | 0.25 | NA | ND | |
| EnvE (52-59) | RPAEVRKV | 0.25 | NA | ND | |
| NS4A (47-56) | MPEAMTIVML | 0.30 | NA | 0/18 | Yes[3] |
| NS3 (588-596) | APGGAKKPL | 0.30 | 13.5 | 0/18 | |
| NS3 (308-318) | RARANESATIL | 0.30 | 3.7 | 0/18 | |
| NS5 (209-219) | NPLSRNSTHEM | 0.30 | 2.6 | 0/18 | |
| NS4B (84-93) | IPFMKMNISV | 0.30 | NA | ND | |
| NS4B (84-94) | IPFMKMNISVI | 0.30 | NA | ND | |
| NS2A (6-13) | VPFGLVSM | 0.30 | 0.4 | 0/16 | |
| CapsidC (37-44) | RPGPSRGV | 0.30 | NA | ND | |
| EnvE (38-45) | KPSLDISL | 0.40 | NA | ND | |
| NS4A (73-80) | SPKGISRM | 0.40 | 1.4 | 0/18 | Yes[2] |
| NS1 (125-133) | SPGRKNGSF | 0.40 | 2.7 | 0/18 | |
| NS3 (227-236) | APTRVVLSEM | 0.40 | 5.3 | 0/18 | |
| NS1 (143-153) | CPFSNRVWNSF | 0.40 | 1.0 | 0/16 | |
| NS4B (51-58) | SPMLHHWI | 0.40 | NA | ND | |
| NS1 (243-252) | MPRSIGGPVS | 0.40 | NA | ND | |
| NS5 (583-591) | RPAPGGKAY | 0.40 | 6.2 | 0 | |
| CapsidC 37-47) | RPGPSRGVQGF | 0.40 | NA | ND | |
| NS2B (71-78) | SARYDVAL | 0.40 | NA | ND | |
| PrM (83-89)) | RSRRSRRAI | 0.40 | NA | ND | |
| NS2B (92-102) | VPWDQVVMTSL | 0.40 | NA | ND | |
| NS4B (205-214) | GPLIEGNTSL | 0.40 | 1.7 | 0 | |
| EnvE ((52-60) | RPAEVRKVC | 0.40 | NA | ND | |
| NS5 672-681) | RPIDDRFGLA | 0.80 | 3.1 | 8/46 | Yes |
| EnvE (284-292) | RVKLSALTL | 1.50 | 4.8 | 5/41 | Yes |

**Table SIII. "hybrid forward-reverse immunology" vs "reverse immunology" approaches to HLA-B*07:02-restricted T cell epitope discovery.**

NetMHCpan 2.4 was used to identify YFV-derived peptides predicted to bind with a high affinity to HLA-B*07:02 (threshold <0.5% Rank), and a selection of these were tested experimentally for their stability of HLA-B*07:02 binding, their ability to support peptide-HLA-B*07:02 tetramer generation, and their ability to stain CD8+ T cells obtained *ex vivo* from YFV vaccinees.

Note: HFRI suffix means that the peptides with the same suffix (1, 2 or3) were found wiithin the same 15mer peptide

| Proteom position | Protein position | Sequence | Positive donors of 50 tested |
|---|---|---|---|
| YFV_pp9-23 | Capsid C (9-23) | KTLGVNMVRRGVRSL | 3 |
| YFV_pp13-27 | Capsid C (13-27) | VNMVRRGVRSLSNKI | 9 |
| YFV_pp17-31 | Capsid C (17-31) | RRGVRSLSNKIKQKT | 14 |
| YFV_pp49-63 | Capsid C (49-63) | FFFLFNILTGKKITA | 11 |
| YFV_pp53-67 | Capsid C (53-67) | FNILTGKKITAHLKR | 9 |
| YFV_pp57-71 | Capsid C (57-71) | TGKKITAHLKRLWKM | 3 |
| YFV_pp61-75 | Capsid C (61-75) | ITAHLKRLWKMLDPR | 4 |
| YFV_pp65-79 | Capsid C (65-79) | LKRLWKMLDPRQGLA | 6 |
| YFV_pp73-87 | Capsid C (73-87) | DPRQGLAVLRKVKRV | 4 |
| YFV_pp77-91 | Capsid C (77-91) | GLAVLRKVKRVVASL | 4 |
| YFV_pp81-95 | Capsid C (81-95) | LRKVKRVVASLMRGL | 10 |
| YFV_pp85-99 | Capsid C (85-99) | KRVVASLMRGLSSRK | 2 |
| YFV_pp121-135 | prM (1-14) | GVTLVRKNRWLLLNV | 1 |
| YFV_pp125-139 | prM (4-18) | VRKNRWLLLNVTSED | 1 |
| YFV_pp133-147 | prM (12-26) | LNVTSEDLGKTFSVG | 1 |
| YFV_pp201-215 | prM (80-94) | SAGRSRRSRRAIDLP | 1 |
| YFV_pp205-219 | prM (84-98) | SRRSRRAIDLPTHEN | 2 |
| YFV_pp213-227 | prM (92-106) | DLPTHENHGLKTRQE | 1 |
| YFV_pp217-231 | prM (96-110) | HENHGLKTRQEKWMT | 1 |
| YFV_pp221-235 | prM (100-114) | GLKTRQEKWMTGRMG | 1 |
| YFV_pp237-251 | prM (116-130) | RQLQKIERWFVRNPF | 2 |
| YFV_pp245-259 | prM (124-138) | WFVRNPFFAVTALTI | 1 |
| YFV_pp249-263 | prM (128-142KK) | NPFFAVTALTIAYLVKK | 3 |
| YFV_pp253-267 | prM (132-146K) | AVTALTIAYLVGSNMK | 4 |
| YFV_pp257-271 | prM (136-150KK) | KKLTIAYLVGSNMTQRV | 1 |
| YFV_pp265-279 | prM (144-158KK) | SNMTQRVVIALLVLAKK | 1 |
| YFV_pp285-299 | EnvE (1-13) | SAHCIGITDRDFIEG | 1 |
| YFV_pp301-315 | EnvE (16-30) | HGGTWVSATLEQDKC | 2 |
| YFV_pp329-343 | EnvE (44-58) | SLETVAIDRPAEVRK | 22 |
| YFV_pp333-347 | EnvE (48-62) | VAIDRPAEVRKVCYN | 5 |
| YFV_pp341-355 | EnvE (56-70) | VRKVCYNAVLTHVKI | 6 |
| YFV_pp345-359 | EnvE (60-74) | CYNAVLTHVKINDKC | 4 |
| YFV_pp409-423 | EnvE (124-138) | SMSLFEVDQTKIQYV | 4 |
| YFV_pp413-427 | EnvE (128-142) | FEVDQTKIQYVIRAQ | 6 |
| YFV_pp417-431 | EnvE (132-146) | QTKIQYVIRAQLHVG | 4 |
| YFV_pp421-435 | EnvE (136-150) | QYVIRAQLHVGAKQE | 9 |
| YFV_pp489-503 | EnvE (204-218) | IVDRQWAQDLTLPWQ | 2 |
| YFV_pp493-507 | EnvE (208-222) | QWAQDLTLPWQSGSG | 1 |
| YFV_pp505-519 | EnvE (220-234) | GSGGVWREMHHLVEF | 2 |
| YFV_pp509-523 | EnvE (224-238) | VWREMHHLVEFEPPH | 3 |
| YFV_pp525-539 | EnvE (240-254) | ATIRVLALGNQEGSL | 3 |
| YFV_pp537-551 | EnvE (252-266) | GSLKTALTGAMRVTK | 1 |
| YFV_pp553-567 | EnvE (268-282) | TNDNNLYKLHGGHVS | 1 |
| YFV_pp557-571 | EnvE (272-286) | NLYKLHGGHVSCRVK | 9 |
| YFV_pp573-587 | EnvE (288-302) | SALTLKGTSYKICTD | 2 |
| YFV_pp589-603 | EnvE (304-318) | MFFVKNPTDTGHGTV | 4 |

| | | | |
|---|---|---|---|
| YFV_pp601-615 | EnvE (316-330) | GTVVMQVKVSKGAPC | 4 |
| YFV_pp613-627 | EnvE (328-342) | APCRIPVIVADDLTA | 1 |
| YFV_pp625-639 | EnvE (340-354) | LTAAINKGILVTVNP | 3 |
| YFV_pp629-643 | EnvE (344-358) | INKGILVTVNPIAST | 1 |
| YFV_pp669-683 | EnvE (384-398) | RLTYQWHKEGSSIGK | 2 |
| YFV_pp673-687 | EnvE (388-402) | QWHKEGSSIGKLFTQ | 6 |
| YFV_pp681-695 | EnvE (396-410) | IGKLFTQTMKGVERL | 4 |
| YFV_pp733-747 | EnvE (KK448-462) | KKGLNWITKVIMGAVLI | 1 |
| YFV_pp745-759 | EnvE (460-474K) | VLIWVGINTRNMTMSK | 3 |
| YFV_pp809-823 | NS1 (31-45) | KYSYYPEDPVKLASI | 3 |
| YFV_pp841-855 | NS1 (63-77) | LEHEMWRSRADEINA | 3 |
| YFV_pp845-859 | NS1 (67-81) | MWRSRADEINAIFEE | 4 |
| YFV_pp849-863 | NS1 (71-85) | RADEINAIFEENEVD | 4 |
| YFV_pp865-879 | NS1 (87-101) | SVVVQDPKNVYQRGT | 7 |
| YFV_pp889-903 | NS1 (111-125) | LQYGWKTWGKNLVFS | 4 |
| YFV_pp893-907 | NS1 (115-129) | WKTWGKNLVFSPGRK | 2 |
| YFV_pp905-919 | NS1 (127-140) | GRKNGSFIIDGKSRK | 2 |
| YFV_pp909-923 | NS1 (131-144) | GSFIIDGKSRKECPF | 4 |
| YFV_pp921-935 | NS1 (143-157) | CPFSNRVWNSFQIEE | 1 |
| YFV_pp945-959 | NS1 (167-181) | VYMDAVFEYTIDCDG | 2 |
| YFV_pp953-967 | NS1 (175-189) | YTIDCDGSILGAAVND | 2 |
| YFV_pp985-999 | NS1 (207-221) | VNGTWMIHTLEALDY | 5 |
| YFV_pp989-1003 | NS1 (211-225) | WMIHTLEALDYKECE | 3 |
| YFV_pp1001-1015 | NS1 (223-237) | ECEWPLTHTIGTSVE | 2 |
| YFV_pp1005-1019 | NS1 (227-241) | PLTHTIGTSVEESEM | 1 |
| YFV_pp1085-1099 | NS1 (307-321) | VIPEWCCRSCTMPPV | 1 |
| YFV_pp1105-1119 | NS1 (327-341) | DGCWYPMEIRPRKTH | 4 |
| YFV_pp1129-1143 | NS2A (1-13) | TAGEIHAVPFGLVSM | 1 |
| YFV_pp1133-1147 | NS2A (3-17KK) | IHAVPFGLVSMMIAMKK | 1 |
| YFV_pp1141-1155 | NS2A (11-25) | VSMMIAMEVVLRKRQ | 1 |
| YFV_pp1177-1191 | NS2A (47-61KK) | KKTLLDLLKLTVAVGLH | 4 |
| YFV_pp1181-1195 | NS2A (51-65) | LLKLTVAVGLHFHEM | 1 |
| YFV_pp1197-1211 | NS2A (KK67-80) | KKNGGDAMYMALIAAFS | 1 |
| YFV_pp1217-1231 | NS2A (87-101) | LIGFGLRTLWSPRER | 3 |
| YFV_pp1229-1243 | NS2A (99-113) | RERLVLTLGAAMVEI | 1 |
| YFV_pp1233-1247 | NS2A (103-117KK) | VLTLGAAMVEIALGGKK | 1 |
| YFV_pp1245-1259 | NS2A (115-129) | LGGVMGGLWKYLNAV | 1 |
| YFV_pp1249-1263 | NS2A (119-133) | MGGLWKYLNAVSLCI | 7 |
| YFV_pp1253-1267 | NS2A (123-137KK) | KKWKYLNAVSLCILTIN | 4 |
| YFV_pp1337-1351 | NS2A (207-221) | QPFLGLCAFLATRIFK | 1 |
| YFV_pp1373-1387 | NS2B (19-33) | GLAFQEMENFLGPIA | 1 |
| YFV_pp1377-1391 | NS2B (23-37) | QEMENFLGPIAVGGL | 1 |
| YFV_pp1385-1399 | NS2B (31-45KK) | KKPIAVGGLLMMLVSVA | 1 |
| YFV_pp1485-1499 | NS3 (1-15) | SGDVLWDIPTPKIIE | 2 |
| YFV_pp1489-1503 | NS3 (5-19) | LWDIPTPKIIEECEH | 1 |
| YFV_pp1505-1519 | NS3 (21-35) | EDGIYGIFQSTFLGA | 2 |
| YFV_pp1529-1543 | NS3 (45-59) | GGVFHTMWHVTRGAF | 1 |
| YFV_pp1533-1547 | NS3 (49-63) | HTMWHVTRGAFLVRN | 11 |
| YFV_pp1537-1551 | NS3 (53-67) | HVTRGAFLVRNGKKL | 1 |

| | | | |
|---|---|---|---|
| YFV_pp1541-1555 | NS3 (57-71) | GAFLVRNGKKLIPSW | 1 |
| YFV_pp1581-1595 | NS3 (97-111) | VQLIAAVPGKNVVNV | 1 |
| YFV_pp1629-1643 | NS3 (145-159) | RNGEVIGLYGNGILV | 2 |
| YFV_pp1633-1647 | NS3 (149-163) | VIGLYGNGILVGDNS | 6 |
| YFV_pp1641-1655 | NS3 (157-171) | ILVGDNSFVSAISQT | 1 |
| YFV_pp1685-1699 | NS3 (201-215) | GAGKTRRFLPQILAE | 4 |
| YFV_pp1689-1703 | NS3 (205-219) | TRRFLPQILAECARR | 5 |
| YFV_pp1693-1707 | NS3 (209-223) | LPQILAECARRRLRT | 8 |
| YFV_pp1697-1711 | NS3 (213-227) | LAECARRRLRTLVLA | 1 |
| YFV_pp1701-1715 | NS3 (217-231) | ARRRLRTLVLAPTRV | 1 |
| YFV_pp1713-1727 | NS3 (229-243) | TRVVLSEMKEAFHGL | 1 |
| YFV_pp1737-1751 | NS3 (253-267) | SAHGSGREVIDAMCH | 1 |
| YFV_pp1753-1767 | NS3 (269-283) | TLTYRMLEPTRVVNW | 4 |
| YFV_pp1765-1779 | NS3 (281-295) | VNWEVIIMDEAHFLD | 15 |
| YFV_pp1769-1783 | NS3 (285-299) | VIIMDEAHFLDPASI | 10 |
| YFV_pp1773-1787 | NS3 (289-303) | DEAHFLDPASIAARG | 1 |
| YFV_pp1777-1791 | NS3 (293-307) | FLDPASIAARGWAAH | 1 |
| YFV_pp1797-1811 | NS3 (313-327) | ESATILMTATPPGTS | 1 |
| YFV_pp1821-1835 | NS3 (337-351) | IEDVQTDIPSEPWNT | 3 |
| YFV_pp1833-1847 | NS3 (349-363) | WNTGHDWILADKRPT | 3 |
| YFV_pp1837-1851 | NS3 (353-367) | HDWILADKRPTAWFL | 16 |
| YFV_pp1841-1855 | NS3 (357-371) | LADKRPTAWFLPSIR | 2 |
| YFV_pp1845-1859 | NS3 (361-375) | RPTAWFLPSIRAANV | 2 |
| YFV_pp1849-1863 | NS3 (365-379) | WFLPSIRAANVMAAS | 10 |
| YFV_pp1853-1867 | NS3 (369-383) | SIRAANVMAASLRKA | 8 |
| YFV_pp1861-1875 | NS3 (377-391) | AASLRKAGKSVVVLNK | 1 |
| YFV_pp1869-1883 | NS3 (385-399) | KSVVVLNRKTFEREY | 4 |
| YFV_pp1877-1891 | NS3 (393-407) | KTFEREYPTIKQKKP | 4 |
| YFV_pp1897-1911 | NS3 (413-427) | TDIAEMGANLCVERV | 1 |
| YFV_pp1929-1943 | NS3 (445-459) | VAIKGPLRISASSAA | 1 |
| YFV_pp1933-1947 | NS3 (449-463) | GPLRISASSAAQRRG | 3 |
| YFV_pp1941-1955 | NS3 (457-471) | SAAQRRGRIGRNPNR | 2 |
| YFV_pp1945-1959 | NS3 (461-475) | RRGRIGRNPNRDGDS | 2 |
| YFV_pp1957-1971 | NS3 (473-487) | GDSYYYSEPTSENNA | 4 |
| YFV_pp1961-1975 | NS3 (477-491) | YYSEPTSENNAHHVC | 2 |
| YFV_pp1969-1983 | NS3 (485-499) | NNAHHVCWLEASMLL | 2 |
| YFV_pp1973-1987 | NS3 (489-503) | HVCWLEASMLLDNME | 1 |
| YFV_pp1989-2003 | NS3 (505-519) | RGGMVAPLYGVEGTK | 1 |
| YFV_pp2021-2035 | NS3 (537-551) | FRELVRNCDLPVWLS | 1 |
| YFV_pp2029-2043 | NS3 (545-559) | DLPVWLSWQVAKAGL | 2 |
| YFV_pp2033-2047 | NS3 (549-563) | WLSWQVAKAGLKTND | 1 |
| YFV_pp2185-2199 | NS4A (78-92) | SRMSMAMGTMAGCGY | 1 |
| YFV_pp2189-2203 | NS4A (82-96K) | MAMGTMAGCGYLMFLK | 2 |
| YFV_pp2285-2299 | NS4B (29-43) | WPDLDLKPGAAWTVY | 1 |
| YFV_pp2289-2303 | NS4B (33-47) | DLKPGAAWTVYVGIV | 3 |
| YFV_pp2293-2307 | NS4B (37-51K) | GAAWTVYVGIVTMLSK | 1 |
| YFV_pp2297-2311 | NS4B (41-55K) | TVYVGIVTMLSPMLHK | 5 |
| YFV_pp2329-2343 | NS4B (73-87) | SASVLSFMDKGIPFM | 1 |
| YFV_pp2337-2351 | NS4B (81-95K) | DKGIPFMKMNISVIMK | 4 |

| | | | |
|---|---|---|---|
| YFV_pp2341-2355 | NS4B (85-99KK) | KKPFMKMNISVIMLLVS | 1 |
| YFV_pp2353-2367 | NS4B (97-111) | LVSGWNSITVMPLLC | 4 |
| YFV_pp2357-2371 | NS4B (101-115KK) | KKWNSITVMPLLCGIGC | 1 |
| YFV_pp2381-2395 | NS4B (125-139) | PGIKAQQSKLAQRRV | 2 |
| YFV_pp2385-2399 | NS4B (129-143) | AQQSKLAQRRVFHGV | 1 |
| YFV_pp2389-2403 | NS4B (133-147 (E->K) | KLAQRRVFHGVAKNP | 5 |
| YFV_pp2393-2407 | NS4B (137-151 (E->K) | RRVFHGVAKNPVVDG | 4 |
| YFV_pp2397-2411 | NS4B (141-155 (E->K) | HGVAKNPVVDGNPTV | 1 |
| YFV_pp2401-2415 | NS4B (146-159) | KNPVVDGNPTVDIEE | 2 |
| YFV_pp2405-2419 | NS4B (149-163) | VDGNPTVDIEEAPEM | 1 |
| YFV_pp2425-2439 | NS4B (KK169-183) | KKKKLALYLLLALSLAS | 1 |
| YFV_pp2437-2451 | NS4B (181-195) | LASVAMCRTPFSLAE | 6 |
| YFV_pp2481-2495 | NS4B (225-239) | TGVMRGNHYAFVGVM | 1 |
| YFV_pp2485-2499 | NS4B (229-241) | RGNHYAFVGVMYNLW | 1 |
| YFV_pp2489-2503 | NS4B (233-247K) | YAFVGVMYNLWKMKTK | 10 |
| YFV_pp2513-2527 | NS5 (7-21) | TLGEVWKRELNLLDK | 2 |
| YFV_pp2517-2531 | NS5 (11-25) | VWKRELNLLDKRQFE | 1 |
| YFV_pp2529-2543 | NS5 (23-37) | QFELYKRTDIVEVDR | 4 |
| YFV_pp2565-2579 | NS5 (59-73) | TAKLRWFHERGYVKL | 2 |
| YFV_pp2569-2583 | NS5 (63-77) | RWFHERGYVKLEGRV | 1 |
| YFV_pp2573-2587 | NS5 (67-81) | ERGYVKLEGRVIDLG | 2 |
| YFV_pp2677-2691 | NS5 (171-185K) | WLACGVDNFCVKVLAK | 3 |
| YFV_pp2833-2847 | NS5 (327-341) | KILTYPWDRIEEVTR | 1 |
| YFV_pp2877-2891 | NS5 (371-385) | TRKIMKVVNRWLFRH | 11 |
| YFV_pp2881-2895 | NS5 (375-389) | MKVVNRWLFRHLARE | 3 |
| YFV_pp2885-2899 | NS5 (379-393) | NRWLFRHLAREKNPR | 3 |
| YFV_pp2905-2919 | NS5 (399-413) | EFIAKVRSHAAIGAY | 1 |
| YFV_pp2957-2971 | NS5 (451-465) | CVYNMMGKREKKLSE | 1 |
| YFV_pp2977-2991 | NS5 (471-485) | GSRAIWYMWLGARYL | 10 |
| YFV_pp2981-2995 | NS5 (475-489(KK) | IWYMWLGARYLEFEAKK | 8 |
| YFV_pp2989-3003 | NS5 (483-497) | RYLEFEALGFLNEDH | 1 |
| YFV_pp3057-3071 | NS5 (551-565) | EQEILNYMSPHHKKL | 5 |
| YFV_pp3061-3075 | NS5 (555-569) | LNYMSPHHKKLAQAV | 2 |
| YFV_pp3069-3083 | NS5 (563-577) | KKLAQAVMEMTYKNK | 1 |
| YFV_pp3073-3087 | NS5 (567-581) | QAVMEMTYKNKVVKV | 2 |
| YFV_pp3077-3091 | NS5 (571-585) | EMTYKNKVVKVLRPA | 2 |
| YFV_pp3117-3131 | NS5 (611-625) | LNTITNLKVQLIRMA | 3 |
| YFV_pp3145-3159 | NS5 (639-653(KK) | KKCDESVLTRLEAWLTE | 1 |
| YFV_pp3205-3219 | NS5 (699-713) | QPSKGWNDWENVPFC | 1 |
| YFV_pp3213-3227 | NS5 (707-727) | WENVPFCSHHFHELQ | 1 |
| YFV_pp3277-3291 | NS5 (771-785) | FHKRDMRLLSLAVSS | 3 |
| YFV_pp3281-3295 | NS5 (775-789) | DMRLLSLAVSSAVPT | 4 |
| YFV_pp3313-3327 | NS5 (807-821) | WMTTEDMLEVWNRVW | 1 |
| YFV_pp3369-3383 | NS5 (863-877) | WASHIHLVIHRIRTL | 2 |
| YFV_pp3373-3387 | NS5 (867-881) | IHLVIHRIRTLIGQE | 13 |
| YFV_pp3377-3391 | NS5 (871-885) | IHRIRTLIGQEKYTD | 9 |

Supplementary Table IV: List of 192 $CD4^+$ T cell stimulatory 15mer epitopes

| Proteome position | Protein position | Sequence | TMR validated HLA-II restriction | Predicted affinity (%RANK) | Measured affinity (nM) | Responders /donors tested |
|---|---|---|---|---|---|---|
| YFV_pp17-31 | Capsid C (17-31) | RRGVRSLSNKIKQKT | DRB5*01:01 | 0.50 | 4 | 6/6 |
| YFV_pp49-63 | Capsid C (49-63) | FFFLFNILTGKKITA | DRB1*01:01 | 0.70 | 27 | 8/8 |
| YFV_pp53-67 | Capsid C (53-67) | FNILTGKKITAHLKR | DRB1*01:01 | 6.50 | 7 | 8/8 |
| YFV_pp65-79 | Capsid C (65-79) | LKRLWKMLDPRQGLA | DRB1*01:01 | 4.50 | 15 | 5/5 |
| YFV_pp73-87 | Capsid C (73-87) | DPRQGLAVLRKVKRV | DRB1*13:01 | 0.12 | 4 | 1/1 |
| YFV_pp73-87 | Capsid C (73-87) | DPRQGLAVLRKVKRV | DRB1*13:02 | 32.00 | 5 | 5/5 |
| YFV_pp77-91 | Capsid C (77-91) | GLAVLRKVKRVVASL | DRB1*14:54 | 0.17 | 6 | 1/1 |
| YFV_pp81-95 | Capsid C (81-95) | LRKVKRVVASLMRGL | DRB1*07:01 | 0.70 | 7 | 5/5 |
| YFV_pp81-95 | Capsid C (81-95) | LRKVKRVVASLMRGL | DRB1*11:01 | 0.25 | 4 | 2/2 |
| YFV_pp85-99 | Capsid C (85-99) | KRVVASLMRGLSSRK | DRB1*11:01 | 0.12 | 5 | 1/1 |
| YFV_pp237-251 | prM (116-130) | RQLQKIERWFVRNPF | DRB1*14:54 | 3.50 | 8 | 1/1 |
| YFV_pp329-343 | EnvE (44-58) | SLETVAIDRPAEVRK | DRB1*03:01 | 0.40 | 21 | 14/15 |
| YFV_pp329-343 | EnvE (44-58) | SLETVAIDRPAEVRK | DRB3*03:01 | 9.00 | 21 | 11/11 |
| YFV_pp333-347 | EnvE (48-62) | VAIDRPAEVRKVCYN | DRB3*03:01 | 32.00 | 12 | 5/7 |
| YFV_pp333-347 | EnvE (48-62) | VAIDRPAEVRKVCYN | DRB1*03:01 | 11.00 | 15 | 4/5 |
| YFV_pp341-355 | EnvE (56-70) | VRKVCYNAVLTHVKI | DRB5*01:01 | 1.30 | 4 | 6/6 |
| YFV_pp345-359 | EnvE (60-74) | CYNAVLTHVKINDKC | DRB5*01:01 | 7.00 | 9 | 5/5 |
| YFV_pp417-431 | EnvE (132-146) | QTKIQYVIRAQLHVG | DRB1*14:54 | 1.90 | 23 | 1/1 |
| YFV_pp525-539 | EnvE (240-254) | ATIRVLALGNQEGSL | DRB1*04:04 | 4.50 | 22 | 2/2 |
| YFV_pp557-571 | EnvE (272-286) | NLYKLHGGHVSCRVK | DRB1*13:02 | 44.00 | 9 | 7/7 |
| YFV_pp673-687 | EnvE (288-402) | QWHKEGSSIGKLFTQ | DRB1*04:01 | 43.00 | 567 | 2/2 |
| YFV_pp601-615 | EnvE (316-330) | GTVVMQVKVSKGAPC | DRB1*04:04 | 13.00 | 16 | 1/1 |
| YFV_pp625-639 | EnvE (340-354) | LTAAINKGILVTVNP | DRB3*03:01 | 1.50 | 2 | 5/5 |
| YFV_pp865-879 | NS1 (87-101) | SVVVQDPKNVYQRGT | DRB1*03:01 | 0.80 | 10 | 5/5 |
| YFV_pp889-903 | NS1 (111-125) | LQYGWKTWGKNLVFS | DRB1*13:01 | 6.50 | 6 | 1/1 |
| YFV_pp889-903 | NS1 (111-125) | LQYGWKTWGKNLVFS | DRB1*13:02 | 30.00 | 9 | 4/4 |
| YFV_pp893-907 | NS1 (115-129) | WKTWGKNLVFSPGRK | DRB1*13:02 | 34.00 | 5 | 3/3 |
| YFV_pp1105-1119 | NS1 (327-341) | DGCWYPMEIRPRKTH | DRB5*01:01 | 1.50 | 2 | 8/8 |
| YFV_pp1373-1387 | NS2B (19-33) | GLAFQEMENFLGPIA | DRB1*15:01 | 16.00 | 42 | 1/1 |
| YFV_pp1533-1547 | NS3 (49-63) | HTMWHVTRGAFLVRN | DRB1*07:01 | 0.01 | 3 | 8/8 |
| YFV_pp1541-1555 | NS3 (57-71) | GAFLVRNGKKLIPSW | DRB1*13:01 | 0.06 | 2 | 1/1 |
| YFV_pp1541-1555 | NS3 (57-71) | GAFLVRNGKKLIPSW | DRB1*13:02 | 1.70 | 3 | 1/1 |
| YFV_pp1541-1555 | NS3 (57-71) | GAFLVRNGKKLIPSW | DRB1*14:54 | 2.50 | 6 | 1/1 |
| YFV_pp1633-1647 | NS3 (149-163) | VIGLYGNGILVGDNS | DRB1*15:01 | 18.00 | 14 | 4/4 |
| YFV_pp1765-1779 | NS3 (281-295) | VNWEVIIMDEAHFLD | DRB1*03:01 | 0.07 | 43 | 5/5 |
| YFV_pp1769-1783 | NS3 (285-299) | VIIMDEAHFLDPASI | DRB1*03:01 | 0.10 | 13 | 6/6 |
| YFV_pp1765-1779 | NS3 (281-295) | VNWEVIIMDEAHFLD | DRB3*01:01 | 0.02 | 3 | 8/10 |
| YFV_pp1769-1783 | NS3 (285-299) | VIIMDEAHFLDPASI | DRB3*01:01 | 0.08 | 8 | 1/1 |
| YFV_pp1765-1779 | NS3 (281-295) | VNWEVIIMDEAHFLD | DRB3*03:01 | 4.00 | 15 | 4/4 |
| YFV_pp1769-1783 | NS3 (285-299) | VIIMDEAHFLDPASI | DRB3*03:01 | 6.00 | 11 | 4/4 |
| YFV_pp1833-1847 | NS3 (349-363) | WNTGHDWILADKRPT | DRB1*03:01 | 8.50 | 28 | 3/3 |
| YFV_pp1837-1851 | NS3 (353-367) | HDWILADKRPTAWFL | DRB1*03:01 | 0.01 | 2 | 12/12 |
| YFV_pp1849-1863 | NS3 (365-379) | WFLPSIRAANVMAAS | DRB1*04:04 | 0.08 | 4 | 1/1 |
| YFV_pp1853-1867 | NS3 (369-383) | SIRAANVMAASLRKA | DRB1*04:04 | 0.80 | 13 | 1/1 |
| YFV_pp1877-1891 | NS3 (393-407) | KTFEREYPTIKQKKP | DRB5*01:01 | 11.00 | 2 | 5/5 |
| YFV_pp2021-2035 | NS3 (537-551) | FRELVRNCDLPVWLS | DRB3*03:01 | 0.70 | 5 | 1/2 |
| YFV_pp2337-2351 | NS4B (81-95K) | DKGIPFMKMNISVIMK | DRB3*03:01 | 0.15 | 3 | 5/5 |
| YFV_pp2401-2415 | NS4B (146-159) | KNPVVDGNPTVDIEE | DRB3*03:01 | 44.00 | 7 | 5/5 |
| YFV_pp2405-2419 | NS4B (149-163) | VDGNPTVDIEEAPEM | DRB3*03:01 | 90.00 | 22 | 5/5 |
| YFV_pp2437-2451 | NS4B (181-195) | LASVAMCRTPFSLAE | DRB1*04:04 | 0.04 | 2 | 1/1 |
| YFV_pp2489-2503 | NS4B (233-247K) | YAFVGVMYNLWKMKTK | DPA1*01:03-DPB1*04:01 | 5.00 | NA | 4/4 |
| YFV_pp2565-2579 | NS5 (59-73) | TAKLRWFHERGYVKL | DRB1*13:01 | 12.00 | 7 | 1/1 |
| YFV_pp2565-2579 | NS5 (59-73) | TAKLRWFHERGYVKL | DRB1*13:02 | 47.00 | 10 | 1/2 |
| YFV_pp2565-2579 | NS5 (59-73) | TAKLRWFHERGYVKL | DRB5*01:01 | 3.00 | 4 | 4/4 |
| YFV_pp2569-2583 | NS5 (63-77) | RWFHERGYVKLEGRV | DRB5*01:01 | 5.50 | 5 | 1/1 |
| YFV_pp2877-2891 | NS5 (371-385) | TRKIMKVVNRWLFRH | DRB1*15:01 | 0.60 | 7 | 5/5 |
| YFV_pp2905-2919 | NS5 (399-413) | EFIAKVRSHAAIGAY | DRB1*15:01 | 1.20 | 2 | 1/1 |
| YFV_pp2977-2991 | NS5 (471-485) | GSRAIWYMWLGARYL | DRB1*01:01 | 6.50 | 6 | 8/8 |
| YFV_pp2981-2995 | NS5 (475-489KK) | IWYMWLGARYLEFEAKK | DRB1*01:01 | 17.00 | 4 | 8/8 |
| YFV_pp3057-3071 | NS5 (551-565) | EQEILNYMSPHHKKL | DRB1*15:01 | 0.15 | 29 | 5/5 |
| YFV_pp3057-3071 | NS5 (551-565) | EQEILNYMSPHHKKL | DRB5*01:01 | 0.08 | 2 | 5/5 |
| YFV_pp3061-3075 | NS5 (555-569) | LNYMSPHHKKLAQAV | DRB5*01:01 | 0.50 | 2 | 3/3 |
| YFV_pp3373-3387 | NS5 (867-881) | IHLVIHRIRTLIGQE | DRB1*04:04 | 0.10 | 14 | 2/2 |
| YFV_pp3377-3391 | NS5 (871-885) | IHRIRTLIGQEKYTD | DRB1*04:04 | 1.80 | 27 | 2/2 |

**Table SV. A complete list of the 50 CD4$^{+}$ T cell epitopes identified by the HFRI approach.**

The CD4$^{+}$ T cell epitope discovery process detailed for donor YF1067 was extended to 50 randomely selected, primary YFV vaccinated donors. 50 different CD4$^{+}$ T cell responses restricted by 14 HLA-II allotypes (13 HLA-DR-restricted and one HLA-DP-restricted) were discovered. The peptide-HLA-I affinities were predicted by NetMHCIIpan (given in %Rank) and the binding affinity was measured (given in nM) (the lower the prediction or measurement, the better the affinity). Productively interacting peptide-HLA-II combinations were used to design and generate peptide-HLA class II tetramers. The resulting tetramers were used to stain and analyze expanded CD4+ T cells by flow cytometry gating on CD3+ CD4+ T cells. The prevalence of these responses is given as "the number of responding donors/the number of donors tested".
In the event that two overlapping peptides were identified, the overlapping sequence has been colored red. For solubilization purpose, lysines have in some cases been added to the N- or C-terminal of the peptide (indicated here by underlining these added lysines.
